# Saccades along spatial neural circuit discontinuities

**DOI:** 10.1101/2021.01.16.426954

**Authors:** Tatiana Malevich, Ziad M. Hafed

## Abstract

Saccades are realized by six extraocular muscles that define the final reference frame for eyeball rotations. However, upstream of the nuclei innervating the eye muscles, eye movement commands are represented in two-dimensional retinocentric coordinates, as is the case in the superior colliculus (SC). In such spatial coordinates, the horizontal and vertical visual field meridians, relative to the line of sight, are associated with neural tissue discontinuities due to routing of binocular retinal outputs when forming retinotopic sensory-motor maps. At the level of the SC, a functional discontinuity along the horizontal meridian was additionally discovered, beyond the structural vertical discontinuity associated with hemifield lateralization. How do such neural circuit discontinuities influence purely cardinal saccades? Using thousands of saccades from 3 rhesus macaque monkeys and 14 human subjects, we show how the likelihood of purely horizontal or vertical saccades is infinitesimally small, nulling a discontinuity problem. This does not mean that saccades are sloppy. On the contrary, saccades exhibit remarkable direction and amplitude corrections to account for small initial eye position deviations due to fixational variability: “purely” cardinal saccades can deviate, with an orthogonal component of as little as 0.03 deg, to correct for tiny target position deviations from initial eye position. In humans, probing perceptual target localization additionally revealed that saccades show different biases from perception when targets deviate slightly from purely cardinal directions. These results demonstrate a new functional role for fixational eye movements in visually-guided behavior, and they motivate further neurophysiological investigations of saccade trajectory control in the brainstem.

**New and Noteworthy:** Purely cardinal saccades are often characterized as being straight. We show how a small amount of curvature is inevitable, alleviating an implementational problem of dealing with neural circuit discontinuities in the representations of the visual meridians. The small curvature functionally corrects for minute variability in initial eye position due to fixational eye movements. Saccades are far from sloppy; they deviate by as little as <1% of the total vector size to adjust their landing position.

## Introduction

Saccade generation involves multiple transformations of reference frames in neurons (Crawford and Guitton 1997; Goossens and van Opstal 2012; Klier et al. 2003; Leigh et al. 1997; Sajad et al. 2020). For example, visual targets for saccades are represented by retinocentric sensory maps, and gaze shift commands are initially also specified as desired displacement vectors in retinocentric coordinates (Goldberg and Bruce 1990; Hepp et al. 1993; Moschovakis and Highstein 1994; Robinson 1972; Schiller and Stryker 1972; Schlag-Rey et al. 1989; van Opstal et al. 1991). Such retinocentric coordinates are known to exist in saccade-related sensory-motor structures, like the superior colliculus (SC) (Klier et al. 2001; Lee et al. 1988; Robinson 1972; Sparks 1988; 1986; van Opstal et al. 1991; Wurtz and Albano 1980). Ultimately, at the level of extraocular muscle innervations, saccade commands are decomposed into the reference frames of the six extraocular muscles that rotate the eyeball (Angelaki and Hess 2004; Carpenter 1988; Leigh et al. 2015; Quaia and Optican 1998; Raphan 1998). With large gaze saccades requiring head movements to increase the dynamic range of the gaze shifts, coordinate transformations for saccades also involve neck muscle reference frames (Corneil et al. 2002; 2004; Crawford et al. 2003; Daemi and Crawford 2015; DeSouza et al. 2011; Freedman 2008; Freedman and Sparks 1997; Klier and Crawford 2003; Sajad et al. 2015; van Opstal and Kasap 2019). All of these transformations operate remarkably optimally (Crawford and Guitton 1997; Goossens and van Opstal 2012), maintaining a delicate balance between accurate landing positions, on the one hand, and rapidity of finishing the gaze shifts to minimize visual disruptions by eye movements, on the other.

As a result of the need to minimize visual disruptions by saccades as much as possible, saccades are highly ballistic in nature, reaching very high peak velocities (Carpenter 1988; Fuchs 1967; Yarbus 1967). However, it is now known that such a ballistic nature of saccades is still under precise control (Abrams et al. 1989; Carpenter 1988; Hafed 2016; Harris and Wolpert 2006; 1998). In fact, the well-known monotonic main sequence relationship of saccade peak velocity versus saccade amplitude (Bahill et al. 1975; Baloh et al. 1975; Zuber et al. 1965) is not fixed. Rather, saccade velocity (for a given movement amplitude) can be modulated by several factors, including reward (Chen et al. 2014; Chen et al. 2013; Muhammed et al. 2020; Reppert et al. 2015; Shadmehr et al. 2019; Takikawa et al. 2002) and arousal (Di Stasi et al. 2011; Di Stasi et al. 2010; Grace et al. 2010; Hopfenbeck et al. 1995; Jurgens et al. 1981; Reilly et al. 2008; Rothenberg and Selkoe 1981). More interestingly, sudden visual flashes resulting in intra-saccadic visual bursts in sensory-motor structures like the SC can be associated with altered saccade kinematics (Buonocore et al. 2017; Buonocore et al. 2016; Buonocore et al. 2020). This suggests precise intra-saccadic control of eye trajectory, and this idea is also supported by suggestions that individual saccades can exhibit trajectory oscillations as they proceed (Ghasia and Shaikh 2014; Ramat et al. 2005; Ramat et al. 2008; Yee et al. 1994).

As part of such precise control, an interesting question emerges with respect to the stage of representing saccade vectors as desired displacement commands. For example, in the SC, it is thought that the locus of activity on the SC map defines the desired saccade amplitude and direction (Cynader and Berman 1972; Lee et al. 1988; Robinson 1972; van Opstal et al. 1991; Wurtz and Goldberg 1972a; b; 1971). Thus, the anatomical location of a neuron dictates the vector that it represents, resulting in a topographic map (Chen et al. 2019; Cynader and Berman 1972; Hafed and Chen 2016; Robinson 1972). However, due to lateralization, there is a structural discontinuity in this map, with the visual field being divided along the vertical meridian into a left and right SC representation. Similarly, it was recently found that the SC also contains a functional discontinuity along the horizontal meridian, with neurons representing the upper and lower visual fields exhibiting significantly different response characteristics (Hafed and Chen 2016). Even in visual cortices, the right, left, upper, and lower visual field quadrants are represented in segregated neural tissue (Daniel and Whitteridge 1961; Van Essen et al. 1984; Van Essen et al. 1986), by virtue of optic nerve and optic tract routing, raising the question of how purely cardinal saccades can be implemented.

Historically, it was suggested that representations near the vertical meridian in the SC (and other sensory and sensory-motor structures) are bilateral, thus allowing coding of purely vertical saccades (Baizer et al. 1991; Dow et al. 1981; Goldberg and Wurtz 1972; Hafed and Chen 2016; Hafed et al. 2009; Leicester 1968; Pigarev et al. 2001; Schneider et al. 2004; Updyke 1974). However, there still seems to be a contralateral preference in SC neurons’ response field (RF) hotspot locations, with only the RF boundaries extending to cross into the ipsilateral side (Chen et al. 2019; Cowey and Perry 1980; Goldberg and Wurtz 1972; Hafed and Chen 2016; Updyke 1974). As a result, it may be asked whether there are ever purely vertical saccades (and similarly for horizontal movements) when there may never be purely vertical (or horizontal) RF hotspot locations in structures like the SC.

Here, we hypothesized that purely cardinal saccades may not need to exist at all, consistent with previous behavioral observations (Bahill and Stark 1977; Dodge 1917; Erkelens and Sloot 1995; King et al. 1986; Quaia et al. 2000; Smit and Van Gisbergen 1990; Viviani et al. 1977). Specifically, if gaze shift commands (e.g. in the SC) are represented by neurons with RF extents that cross a meridian (e.g. the vertical meridian) but always with preferred RF hotspot locations on one side of the meridian (e.g. due to lateralization), then it may be that purely cardinal saccades are not possible. More importantly, it may be the case that such purely cardinal movements are unnecessary in the first place. Specifically, to get a purely cardinal saccade, the initial eye position at saccade onset needs to be perfectly aligned (along the cardinal axis) with the appearing saccade target location. However, with constant variability in eye position during fixation (Barlow 1952; Steinman et al. 1973), the likelihood of such perfect retinotopic alignment is infinitesimally small, and this is even more true when natural behaviors, including small head movements, come into play. As a result, some small deviation in cardinal saccades might be obligatory. We suggest that such deviation should be sensitive to the directional errors introduced by the scale of fixational eye movement variability. This would, in turn, imply that one additional functional role for fixational eye movements in visually-guided behavior is to avoid visual field discontinuities associated with representing purely cardinal saccade vectors in topographically organized neural circuits. Therefore, saccade trajectory variability may not be entirely due to noise; instead, saccades are remarkably highly directionally sensitive to still accurately allow target foveation with variable initial fixational eye position. In what follows, we provide our evidence supporting this hypothesis. In addition, because visual field discontinuities influence not only saccade metrics (Collewijn et al. 1988b; Hafed and Chen 2016; Henriques and Crawford 2000; Laidlaw and Kingstone 2010; Van der Stigchel and Theeuwes 2008; Zhou and King 2002), but also peri-saccadic perception (Grujic et al. 2018) and spatial perception in the absence of saccades (Greenwood et al. 2017; He et al. 1996; Talgar and Carrasco 2002; Uddin et al. 2004), we also relate our eye movement findings to perceptual localization performance in humans.

## Methods

### Ethics approvals

We performed experiments on 3 male rhesus macaque monkeys (A, M, and N), aged 6, 6, and 12 years, respectively. The experiments were approved by ethics committees at the Regierungspräsidium Tübingen, and they were in line with the European Union directives and the German laws governing research on animals.

Our human experiments were performed on 14 adult subjects (seven females) who were aged 20-40 years. The subjects were compensated 10 Euros per session of approximately 60 minutes, and they provided informed consent. The experiments were approved by ethics committees at the Medical Faculty of Tübingen University, and they were in accordance with the Declaration of Helsinki.

### Monkey laboratory setup

The monkey experiments were performed in the same laboratory as that described in (Buonocore et al. 2019; Malevich et al. 2020; Skinner et al. 2019). The setup consisted of the monkeys being seated in front of a CRT display monitor running at 120 Hz, and with a viewing distance of approximately 73 cm. Display resolution was 34 pixels/deg. The heads of the monkeys were surrounded by a field coil system for tracking eye movements (Remmel Labs, USA). The experimental control system consisted of a real-time computer (National Instruments, USA) that was controlling the behavioral tasks and rewards. The same computer was also monitoring animal behavior. For driving graphics, the real-time computer sent display update commands, via UDP packets, to a computer (Apple, USA) running the Psychophysics Toolbox (Brainard 1997; Kleiner et al. 2007; Pelli 1997). The whole system was described in detail earlier (Chen and Hafed 2013; Chen et al. 2015; Tian et al. 2016). Data and events were stored using a multi-channel acquisition processor (MAP) from Plexon.

### Human laboratory setup

The human experiments were performed in the same laboratory as that described recently (Grujic et al. 2018; Idrees et al. 2020). Briefly, subjects sat in front of a CRT display monitor running at 85 Hz. Viewing distance was 57 cm, and the display had a resolution of 41 pixels/deg. Head fixation was achieved through a custom-built chin and forehead rest, which also had a temple guide and a headband (Hafed 2013). Eye movements were tracked using a video-based eye tracker (EyeLink 1000, SR Research Ltd, Canada) placed under the display and aimed at the left eye, and the eye tracker sampled eye positions at 1000 Hz. The experiments were controlled using the Psychophysics Toolbox (Brainard 1997; Kleiner et al. 2007; Pelli 1997), with EyeLink extensions (Cornelissen et al. 2002). Data and events were stored and analyzed with Matlab (MathWorks Ltd, USA).

### Animal preparation

Each monkey was implanted with a head holder for head fixation during the experiments. In a subsequent surgery, an eye coil was implanted in one eye to allow eye tracking with the precise magnetic induction technique (Fuchs and Robinson 1966; Judge et al. 1980).

Monkeys M and N had the coil implanted in the right eye, and monkey A had the coil implanted in the left eye. Our surgery procedures were described earlier (Chen and Hafed 2013; Chen et al. 2015; Willeke et al. 2019). All 3 monkeys also had chamber implants, for physiological experiments that were beyond the scope of the present study.

### Monkey behavioral task

Each trial started with a white fixation spot (86 cd/m^2^ for monkeys M and A and 72 cd/m^2^ for monkey N) presented at the center of a uniform gray background (29.7 cd/m^2^ for monkeys M and A and 21 cd/m^2^ for monkey N). The spot measured approximately 5 by 5 min arc (Willeke et al. 2019). After the monkey fixated the spot by 300-1500 ms, the spot jumped along one of the cardinal directions (right, left, up, or down) by either 5 deg or 10 deg. The orthogonal component of the target jump could be displaced from pure cardinality by either 0, +/- 1, +/- 2, or +/- 3 pixels (i.e. by 0 min arc, +/- 1.76 min arc, +/- 3.53 min arc, or 5.29 min arc). For example, for a rightward target jump of 5 deg and +1 pixel orthogonal offset, the horizontal component of the jump was 5 deg, and the vertical component was 1.76 min arc (0.03 deg) above the horizontal meridian of the display. Similarly, for an upward target jump of 10 deg and +1 pixel orthogonal offset, the vertical component of the jump was 10 deg, and the horizontal component was 1.76 min arc (0.03 deg) to the right of the vertical meridian. Therefore, the direction of the target vector could deviate from a purely cardinal direction by as little as 0.34 deg (angular deviation) for 5 deg target jumps, and by as little as 0.17 deg (angular deviation) for 10 deg target jumps. The task of the monkey was to generate a visually-guided saccade to the new target location, and we analyzed the saccade trajectories. The monkeys were rewarded for making the saccade within 500 ms from target jump and maintaining gaze near the target for another 500 ms.

We collected 4000-8500 trials from each monkey; these were collected over approximately 16-32 sessions (which often had other tasks performed by the monkeys for other projects). The target eccentricity, direction, and offset from the cardinal axis were randomized within each session. We should also note here that the monkeys were all experts in precise oculomotor tasks. For example, monkey N was used extensively for image stabilization experiments controlling foveal eye position error based on precise fixation control (Tian et al. 2018; 2016), and monkeys M and A were well trained to precisely control their eye position even with foveal targets that moved very slowly (Skinner et al. 2019). Examples of how well trained the monkeys were in fixation control can also be seen in (Malevich et al. 2020).

### Human behavioral tasks

The humans performed two behavioral tasks. The first one was similar to the monkey task, and the second one was a fixation control version of it. In both tasks, the humans made a perceptual judgement about target location, in addition to the required eye movement behavior.

In the first task, the subjects fixated a white spot (97.3 cd/m^2^) presented over a gray background (20.5 cd/m^2^). The spot was placed at the center of the display. After 800-1700 ms, the spot jumped to the right, left, up, or down by 7 deg. The orthogonal component of the cardinal target jump could deviate from pure cardinality by 0, +/- 2, +/- 4, or +/- 8 pixels (i.e. by 0 min arc, +/- 2.93 min arc, 5.85 min arc, or 11.71 min arc). For example, for 7 deg rightward and +2 pixels, the horizontal component of the target jump was 7 deg in size, and the vertical component was 0.05 deg upward. Therefore, the target jump vector could deviate, in angular direction, from a purely cardinal direction by as little as 0.35 deg. The instruction to the subjects was to make a saccade to follow the target jump. When we detected the saccade (online) using a velocity criterion (Baumann et al. 2020; Idrees et al. 2020), we removed the saccade target approximately 50 ms after online detection, in order to avoid having a visual reference after saccade landing. Across all sessions and subjects, we re-detected saccades offline during data analysis, and we measured when the target blanking actually happened: target blanking occurred −14.48 ms +/- 9.59 ms s.d. from saccade end. That is, on average, the target was blanked intra-saccadically before saccade end. After the subjects made the saccade, a question was displayed on the blank screen; for horizontal target jumps, the question was “Above horizon or below horizon?”, and for vertical target jumps, the question was “Right of vertical or left of vertical?”. Subjects had to respond, via a button response box, indicating whether they thought the orthogonal component of the target jump was deviated from pure cardinality. Specifically, for horizontal target jumps and saccades, the subjects pressed an upper button on the response box if they thought that the peripheral target was displaced slightly above the horizontal meridian of the display, and they pressed a lower button on the response box if they thought that the peripheral target was displaced slightly below the horizontal meridian. Similarly, for vertical target jumps and saccades, the subjects pressed a right button on the response box if they judged the peripheral target to have appeared slightly to the right of the vertical meridian, and they pressed a left button if they felt that the peripheral target was displaced slightly to the left of the vertical meridian. Therefore, this task was a dual task (eye movement followed by perceptual report), and we called it the “perceptual dual” task in Results.

Because dual tasks can sometimes cause interactions in performance for either vision or eye movements or both (Harrison et al. 2013; Uddin et al. 2004), we also ran our human subjects in a second control task. In the control task, the subjects maintained fixation. The fixation spot never disappeared, but a second target appeared peripherally exactly as in the saccade task. The target was only presented for 100 ms and then removed, simulating the duration of peripheral preview before saccadic reaction times in the main task. The subjects maintained fixation, and only performed the perceptual report component of the task. We, therefore, referred to this task as the “perceptual control” task. In this task, the target could be displaced away from pure cardinality only by 0, +/- 2, or +/- 8 pixels.

We recruited 17 participants for the human component of this study; two participants withdrew from the study, and one subject was excluded later due to bad performance (see *Eye movement analyses*), which resulted in a sample size of 14 subjects. All participants were naïve to the purposes of the study, reported no history of neurological or psychiatric disorders, were right-handed, and had normal or corrected-to-normal vision. For each subject, we collected 2240-2800 trials from the dual task and 700 trials from the control task. Usually, the subject completed the perceptual dual task in five sessions; the perceptual control task was collected in a separate session. For both tasks, the target direction and offset from cardinality were randomized within a session.

### Eye movement analyses

Saccade detection was performed using established methods reported elsewhere (Bellet et al. 2019; Chen and Hafed 2013); after that, all trials were manually inspected to ensure correct detection and removal of blinks and artifacts. We applied this approach to both the scleral search coil (monkeys) and video-based eye tracking (humans) data sets.

For the monkey data, we excluded trials from analysis if eye movements or blinks occurred in the interval of 100 ms before target jump until saccade initiation (monkey N: 34.66%; monkey M: 28.03%; monkey A: 37.13%); if saccadic latency was either less than 50 ms or exceeded 500 ms (monkey N: 0.16%; monkey M: 0.06%; monkey A: 0.00%); if the saccade end point was more than 15% of the target eccentricity away from the target (monkey N: 5.78%; monkey M: 5.46%; monkey A: 9.39%); and if the Euclidian distance of the eye position at saccade initiation was more than 6 median absolute deviations away from the median fixation position (monkey N: 0.82%; monkey M: 0.43%; monkey A: 0.57%). After the exclusion, on average, there were 89.82 +/- 7.26 s.d. (monkey N), 57.27 +/- 6.36 s.d. (monkey M), and 38.25 +/- 7.13 s.d. (monkey A) trials left per condition. This means that we analyzed a total of 5030, 3207, and 2142 trials from monkeys N, M, and A, respectively.

For the human data, we excluded from analysis the first twenty trials of the first session as practice trials. In the dual task, trials were also excluded if there were eye movements or blinks (14.93%) within the interval of 100 ms before the target jump until saccade initiation; if saccadic latency was either less than 70 ms or larger than 500 ms (2.29%); if the error of the eye position at saccade onset exceeded 1 deg (7.92%); if the Euclidian distance of the saccade end point was more than 3 deg away from the target location (4.43%); if the manual reaction time to the perceptual question exceeded 3000 ms (0.07%); and if the target was removed from the screen before the response saccade was initiated (due to erroneous online saccade detection of noisy eye tracker transients; 4.66%). As mentioned in *Human behavioral tasks*, one participant with only 39% of trials left was excluded from further analysis. After the exclusion, on average, there were 66.14 +/-16.46 s.d. trials left per condition per participant. In the perceptual control task, trials were excluded from analysis if there were eye movements or blinks (15.68%) in the interval of +/-100 ms relative to the target jump and if the manual reaction time to the perceptual question exceeded 3000 ms (0.13%). After the exclusion, on average, there were 29.92 +/- 3.49 s.d. trials per participant per condition.

Unless otherwise stated, all of the analyses for the monkey data were performed separately for each monkey, saccade direction, and target eccentricity. First, we aligned eye traces horizontally and vertically to the eye positions at saccade onset (unless the analysis explicitly looked for effects of initial fixational eye position variability as in Figs. 3, 4 in Results). Then, we plotted the time courses of the mean horizontal and vertical eye position traces for each orthogonal offset in the interval of −50 to 100 ms relative to saccade onset, with the standard error of the mean as an estimate of the across-trial variability. As the next step, we repeated this procedure to obtain the two-dimensional visualization of the time-courses of saccade trajectories by plotting the mean horizontal and mean vertical eye position traces against each other, starting from saccade initiation up to the time of the end point of the shortest saccade in the particular condition.

**Figure 1.**
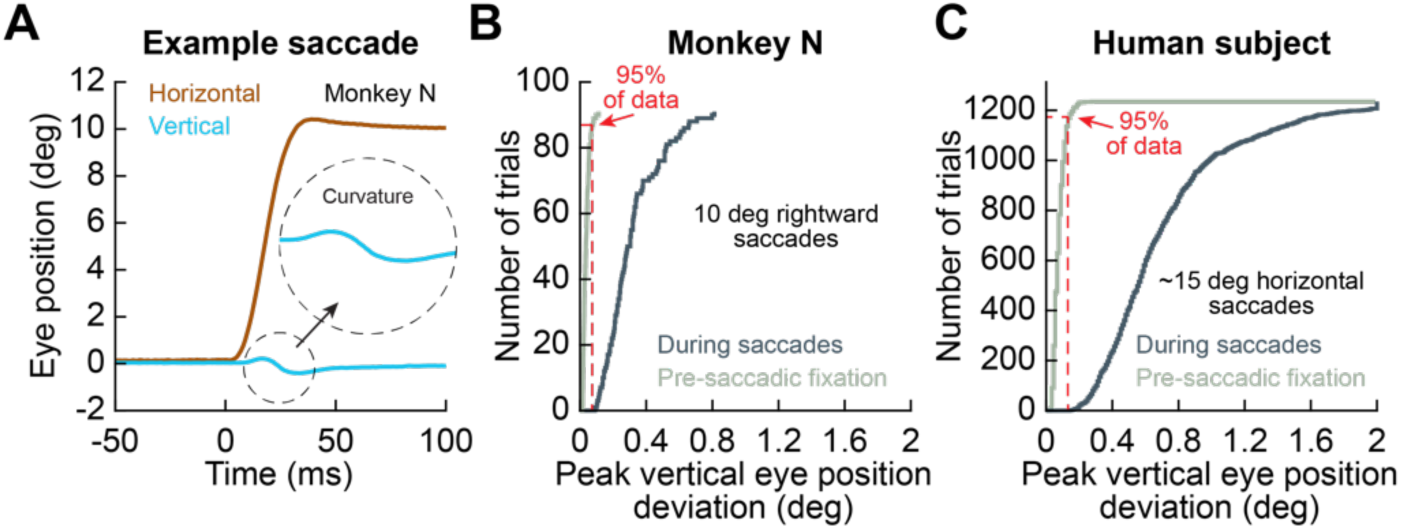
Obligatory deviation from pure cardinality in saccades. **(A)** Horizontal and vertical eye position around an example 10 deg rightward visually-guided saccade in one monkey (eye position was measured using a scleral search coil). The vertical component of eye position during the saccade exhibited a transient deviation, indicating a small amount of curvature, even though the saccade was supposed to be purely horizontal (Bahill and Stark 1977). The peak deviation of vertical eye position during the saccade was significantly larger than the maximal vertical deviation of eye position during the pre-saccadic baseline fixation interval. **(B)** Across tens of repetitions of the same saccade as in **A**, we measured the maximal deviation in vertical eye position during a baseline pre-saccadic fixation interval (light greenish) and during the saccade itself (dark greenish; Methods). The two curves show the cumulative histograms of the measurements across all saccades. Practically every single saccade deviated from being purely horizontal, as indicated by a significantly larger vertical deviation during the saccade than during fixation. The vertical red line shows the extent of vertical eye position deviation from 95% of all of the fixation data, and it overlaps with significantly <1% of the other distribution. **(C)** In an example human subject, we analyzed >1000 repetitions of the same ∼15 deg horizontal saccade (this time, eye position was measured using a video-based eye tracker). The same observations as in the monkey were made. The data for this panel were obtained from the saccades collected in an earlier study (Grujic et al. 2018), but later figures allow similar conclusions with the new data of the current article. The numbers of trials per condition are explicitly shown in the relevant panels (y-axes in **B, C**).

**Figure 2.**
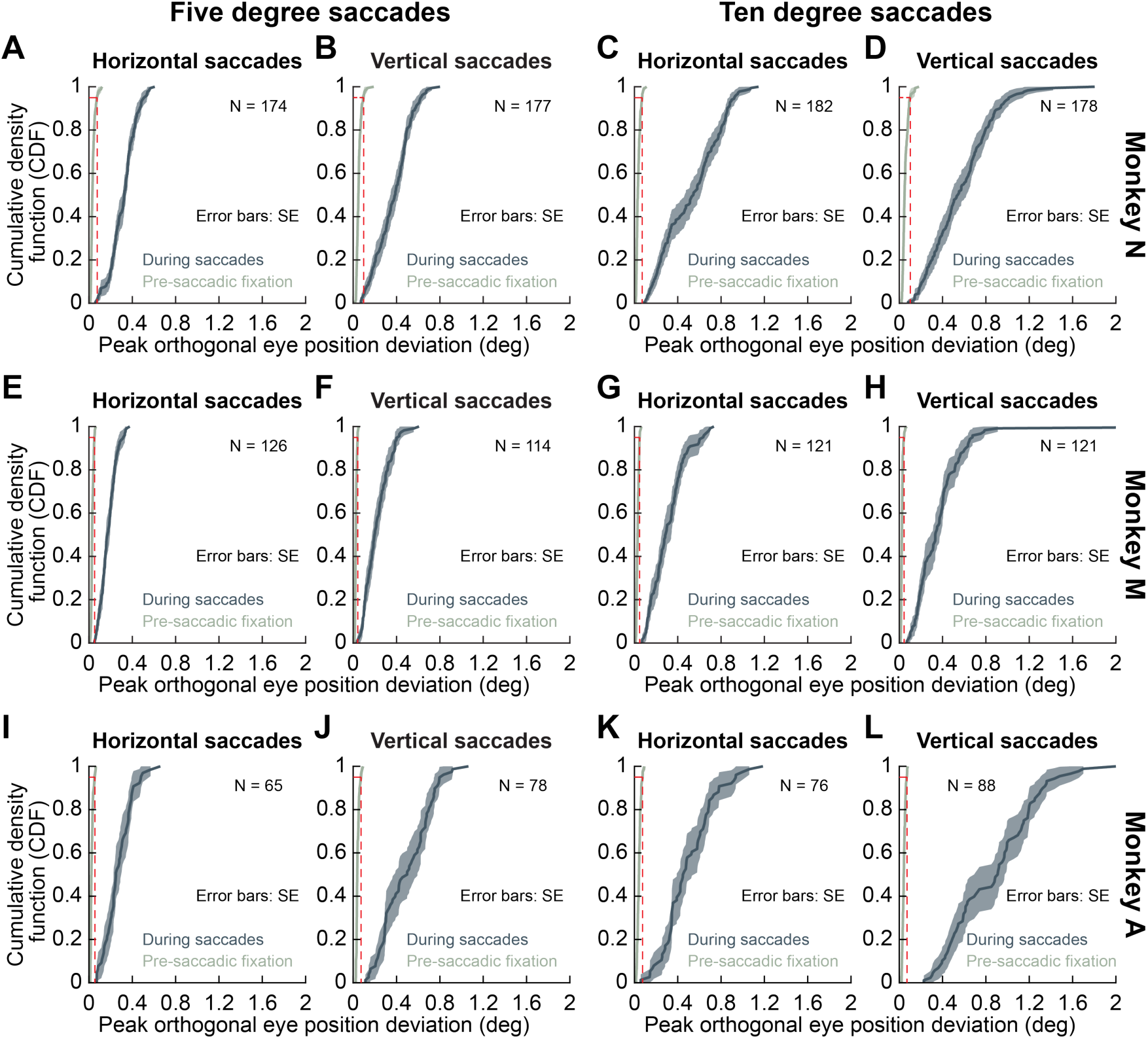
Consistency of the observations of Fig. 1 across different subjects, different saccade sizes, and different cardinal directions. Each row of panels shows results from a single monkey (all tracked using scleral search coils), and each column shows a given saccade condition; the left two columns show results from 5 deg saccades (horizontal and vertical, respectively), and the right two columns show results from 10 deg saccades. In all cases, the target jumped from central fixation by either 5 deg or 10 deg in a purely cardinal direction (right, left, up, or down). Yet, there was always an orthogonal component to the saccades, indicating a minute amount of saccadic curvature. The curvature was also larger with 10 deg saccades than with 5 deg saccades, evidenced by the larger peak deviation along the orthogonal component of the saccade vector. Note that in this figure, for each cardinal direction (e.g. 5 deg horizontal saccades), we combined target jumps of opposite directions (e.g. 5 deg rightward and 5 deg leftward together, or 5 deg upward and 5 deg downward together) in order to simplify the figure. Later figures show separate analyses for each of the four cardinal directions (with consistent results).

**Figure 3.**
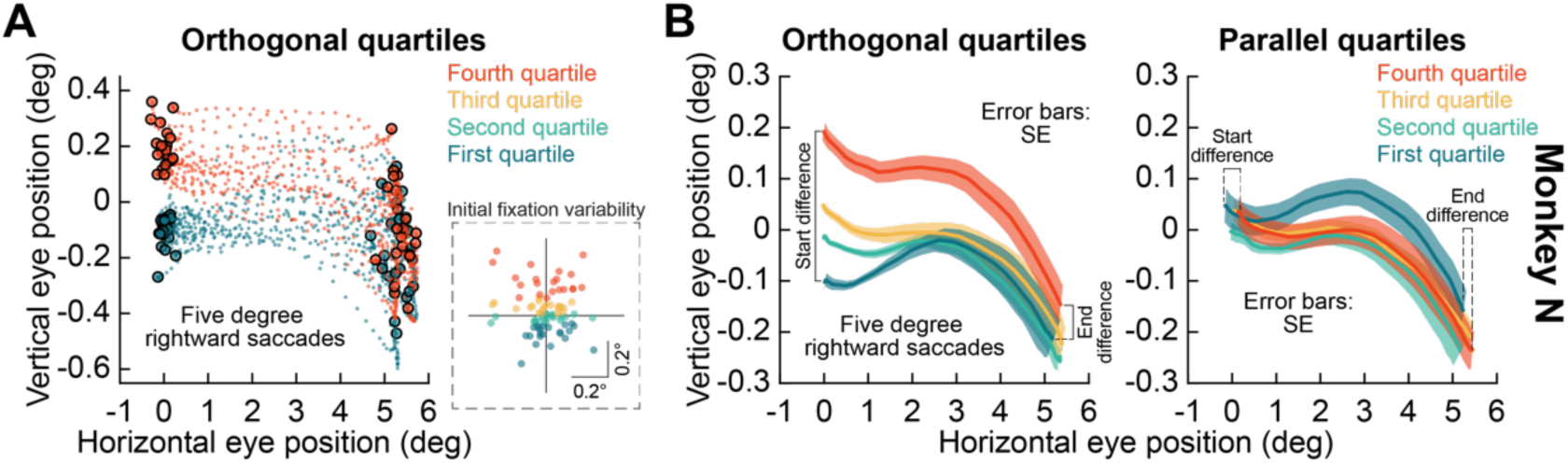
Curvature in cardinal saccades helps to compensate for deviations in initial eye position at saccade onset, due to fixational eye movements. **(A)** For pure 5 deg rightward target jumps in one example monkey, we analyzed saccade trajectories in relation to eye position at saccade onset. Due to fixational eye movements, the saccade (for the same purely rightward target jump) was initiated with different starting eye positions (inset). We divided the trials into four quartiles based on the vertical component (orthogonal to the saccade vector) of initial eye position at saccade onset (four colors in the inset). The main panel shows the trajectories of all saccades from the first and fourth quartiles: intra-saccadic eye position converged by movement end despite the initial divergence between the quartiles at saccade start; thus, a component of saccade curvature in purely cardinal saccades corrects for initial fixational eye position variability. **(B)** Average trajectories from all four quartiles in **A**. The end difference in orthogonal component of eye position (vertical in this case) between the first and fourth quartiles was significantly smaller than the start difference, consistent with **A. (C)** To check whether compensation could also occur in saccade radial size, as opposed to only direction as in **A, B**, we also divided the trials into four quartiles based on horizontal eye position at saccade onset (i.e. parallel to the vector direction). This time, the start difference was the difference in mean horizontal (parallel) eye position at saccade onset between the first and fourth quartiles, and the end difference was the difference in horizontal (parallel) eye position between the same quartiles at saccade end. Remarkably, even the horizontal (parallel) component of the saccades compensated for the initial horizontal eye position variability at saccade onset (see Fig. 4). N = 86 saccades.

**Figure 4.**
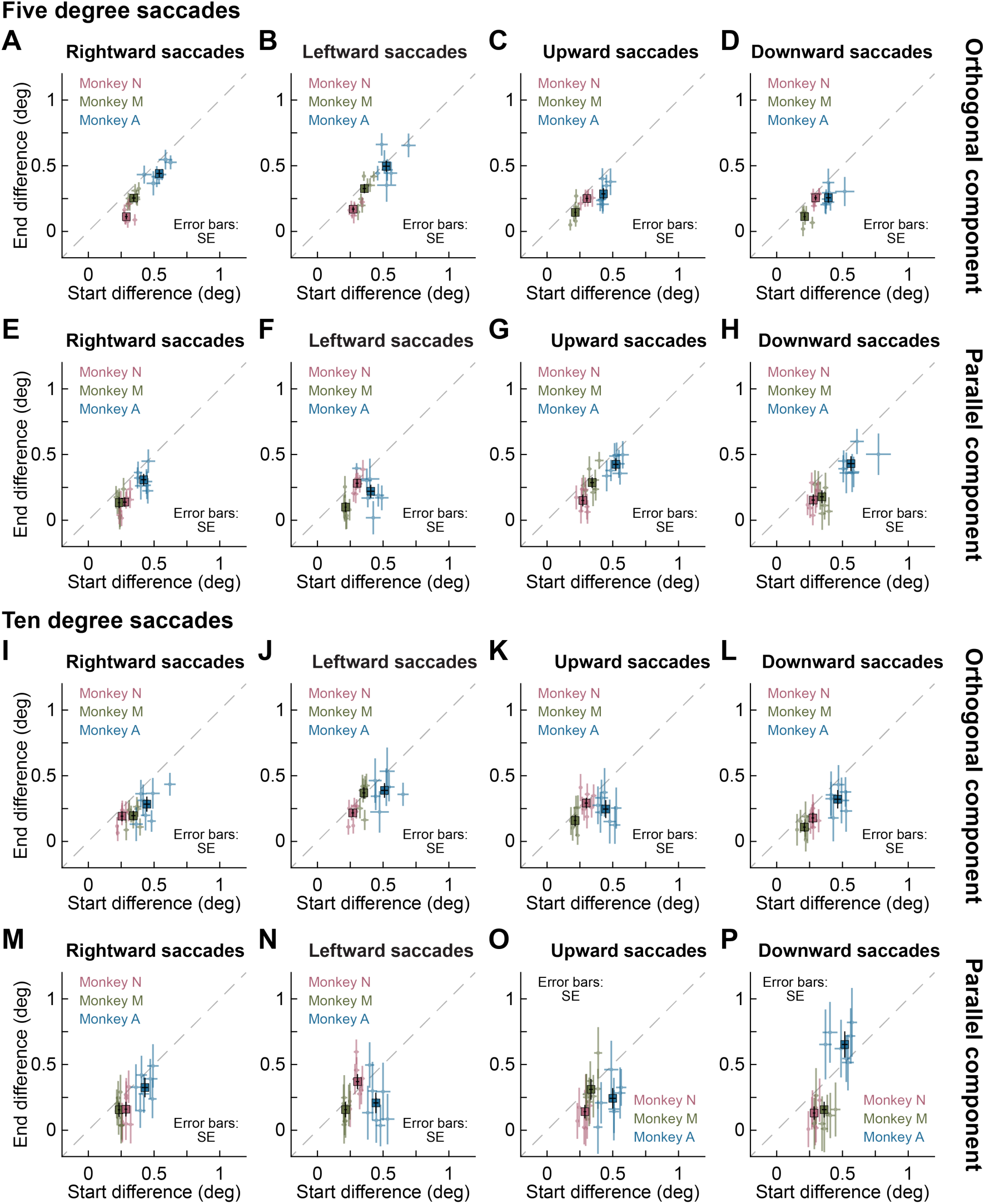
Saccades consistently compensate, in both their direction and amplitude, for initial fixational eye position variability at movement onset. **(A-D)** For each tested cardinal saccade direction (5 deg eccentricity), we plotted the end difference in orthogonal eye position as a function of start difference of orthogonal eye position for each monkey (identical analysis to Fig. 3A, B). Specifically, for each condition, we divided saccades into 4 quartiles based on the orthogonal component of initial eye position at saccade onset. Then, we measured the average (orthogonal) eye position at saccade onset for the first and fourth quartiles; we also measured the average (orthogonal) eye position at saccade end. If saccades compensate in direction for the initial orthogonal fixational eye position variability (Fig. 3A, B), then end difference should be smaller than start difference (Fig. 3A, B). This was true for all monkeys. Each symbol for one monkey shows the analysis for one of the pixel offsets of orthogonal target jump in each condition (Methods). The large symbols show the grand summary across all pixel offset conditions per monkey (Methods). **(E-H)** Same analyses for the parallel component of initial eye position variability (and the size of the parallel component of the saccade), as in Fig. 3C. In all monkeys, even the parallel component size compensated for initial eye position. Thus, a difference of <0.2 deg in parallel eye position at saccade onset was still compensated for in a movement that was >20 times in size (5 deg). N = 70 to 103 saccades per pixel offset and target direction in monkey N, 48 to 71 saccades in monkey M, and 20 to 49 saccades in monkey A. **(I-P)** Same analyses for 10 deg saccades. Similar observations could be made. Note that in this case, parallel corrections meant that a movement >50 times the size of the initial fixational eye position variability (< 0.2 deg) was still compensated in size (monkey A did not show this effect for downward saccades, but this monkey also had the smallest amount of data collected; Methods). N = 77 to 106 saccades per pixel offset and target direction in monkey N, 42 to 72 saccades in monkey M, and 31 to 49 saccades in monkey A.

To investigate saccadic curvature, we first plotted empirical cumulative density functions (empirical CDF’s) of the absolute peak eye position deviation during the saccade itself (for the orthogonal component of intra-saccadic eye position deviation relative to the saccade vector direction) as well as for the absolute peak eye position deviation (again for the orthogonal component) in the final 100 ms before saccade initiation. For example, if the saccade was rightward with a 0 pixel orthogonal target position offset, then the saccade was theoretically “purely rightward” and with no “vertical” component. To identify the vertical curvature of this saccade, we measured the peak intra-saccadic vertical eye position (orthogonal to the saccade vector), and we compared it to the peak vertical eye position deviation during pre-saccadic fixation. From the CDF of the pre-saccadic measurement, we estimated the 95^th^ percentile, and we checked whether this value was still smaller than peak orthogonal intra-saccadic eye position deviation. For the final report (e.g. in Figs. 1, 2), we used only the 0 pixel conditions and collapsed the trials for horizontal (i.e. rightward and leftward) and vertical (i.e. upward and downward) saccades. However, we ensured that the same analysis performed separately on each saccade direction and each pixel offset condition brought analogous results. To directly test the hypothesis that the curvature of larger saccades was larger than the curvature of smaller saccades, we ran a one-sided two-sample Kolmogorov-Smirnov (K-S) test on the absolute peak eye position intra-saccadic orthogonal deviations of five-versus ten-degree saccades.

Further, we compared the initial fixational eye position variability to the variability at the saccade end points and investigated the idea that the former is reduced during the course of a saccade. We, specifically, first asked whether a deviation in initial fixational eye position orthogonal to the direction of the upcoming saccade caused more curvature in the saccade (to correct for the initial eye position error). In order to answer this question, for each pixel offset condition and each saccade direction, we divided the trials into four quartiles based on the orthogonal component of initial eye position at fixation onset. For example, if the saccade was 5 deg rightward and with +1 pixel offset, we divided all trials from this condition based on the vertical (orthogonal) component of eye position at saccade onset. Similarly, if the saccade was 10 deg upward, we divided the trials into four quartiles based on horizontal (orthogonal) eye position at saccade onset. We then plotted the average saccade trajectories for each quartile and their standard errors; here, the data were not aligned to the saccade onset eye position since we were interested in how the saccades compensated for initial variability in fixational eye position. Next, we calculated the differences between the upper and lower quartiles’ mean values at the saccade onset (called “start difference”) and at the end of the saccades (called “end difference”). The ratio of the difference of the quartiles’ means at the saccade end to the difference of the quartiles’ means at the saccade onset gave us the Initial Fixation Variability Compensation Indices (IFVCI’s); IFVCI’s < 1 indicated that there was a correction for the initial variability, whereas IFVCI’s >= 1 showed no correction. In plots, trials with IFVCI’s <1 showed end differences lying below the unity slope line of graphs of end versus start differences (e.g. Fig. 4 in Results). To assess the correction for initial fixation variability irrespective of the orthogonal offset, we performed a bootstrapping procedure; namely, for 1000 times, we randomly resampled our upper and lower quartiles’ data with replacement, computed the IFVCI’s for each orthogonal offset condition, and took their medians. In the end, this procedure provided us with the distribution of 1000 median IFVCI values. The central tendency measure and the estimate of its standard error were calculated as the mean and standard deviation of the bootstrap distribution (e.g. Fig. 4 in Results). Finally, we computed Monte Carlo p-values by assessing the probability of getting bootsrap IFVCI values larger than or equal to 1 (i.e. by taking the count of the bootsrap IFVCI values larger than or equal to 1 and dividing this count by the number of resamples).

We also repeated the same analysis again, but this time dividing initial fixational eye positions into four quartiles based on the parallel component of eye position relative to the saccade direction. That is, for a 5 deg rightward saccade condition, we divided all trials of this condition into four quartiles based on the horizontal (parallel) eye position at saccade onset. This allowed us to ask whether saccades also adjusted their radial amplitudes to account for initial fixational eye position variability bringing the eye ever so slightly closer to or farther away from the peripheral saccade target.

To statistically determine the sensitivity of saccade trajectory (as a function of time) to the orthogonal pixel offsets that we introduced experimentally, we fitted generalized additive mixed models (GAMM’s), a nonparametric regression method, which uses smooth functions (so-called ‘smoothers’ or ‘smooths’) to model nonlinear relationships between the response variable and explanatory variables, and also allows inclusion of nonlinear random effects (Wood 2017; Wood et al. 2015). The analysis was performed in R (version 3.6.1; R Core Team 2019), using packages mgcv (version 1.8-28) (Wood 2017) and itsadug (version 2.3) (van Rij et al. 2015). Due to a low number of trials per condition in monkey A, only monkeys N and M were included in this particular analysis. The models were fitted separately for each monkey, direction, and target eccentricity. For this analysis, we normalized saccades to their onset eye position (as stated earlier in the analyses not related to initial fixational eye position variability). For each model, relevant eye traces were cut off at the time point at which their duration exceeded the 90^th^ percentile of all saccade durations; this was necessary to ensure equal numbers of repetitions across all time samples during the analyzed intra-saccadic periods. The models were then fitted with the bam() function of the mgcv package (Wood 2017). The orthogonal eye position (relative to the saccade direction) was the dependent variable. As fixed effects, the full model included a seven-level categorical predictor of the target pixel offset (i.e. from −3 to +3 pixel offsets) and a factor-smooth interaction of the target pixel offset and time (during a saccade), which produced a separate smoother for each level of the target pixel offset over time. The random part of the model’s structure was modeled as a by-session random smoother of time (during a saccade) (Wieling 2018; Wood 2017) to account for a potential between-session variability in the time courses of saccade trajectories. The best-fit models were determined with a backward-fitting comparison procedure. That is, in each particular case, we started from a full model and compared it to the reduced model that included only random effects and no fixed effects. The models were assessed against each other based on their difference in Akaike’s information criterion (AIC) scores (Akaike 1998). AIC-scores provide the models’ relative goodness of fit based on their likelihood while penalizing a more complex model for the number of free parameters, and a model with the lowest AIC-score is considered to be the best-fit model. This procedure was performed with the compareML() function of the itsadug package (van Rij et al. 2015). When comparing models with different fixed effects, we fitted the models using the maximum likelihood estimation method. To determine the optimal model’s random effect structure, we used a similar procedure but fitted models with the fast restricted maximum likelihood estimation method, which is suited for estimating variance from the random effects in mixed models (Wieling 2018). To account for the presence of autocorrelations in the residuals (i.e. in the differences between the observed values of the dependent variable and the predicted values), we incorporated an autoregressive error model of order 1 for the residuals in the best-fit model’s structure (Baayen et al. 2017; Wood 2017). Finally, we assessed the difference between the smoothers estimated for saccade trajectories of the adjacent orthogonal pixel offsets with the help of the plot_diff() function of the itsadug package (van Rij et al. 2015); in particular, this function uses a simulation-based approach to calculate the area of the surface where the difference curve between a pair of estimated smoothers is different from zero at 95% simultaneous confidence level.

To assess the discriminability of individual saccades as a function of orthogonal pixel offset values in the target jumps, we next followed a procedure similar to the one described in (Yoshida et al. 2008). Namely, we calculated the area under the curve (AUC) of the receiver operating characteristic (ROC) curve from the distribution of the orthogonal components of the saccade end points in the 0 pixel condition and the distribution of the orthogonal components of the saccade end points in a condition, in which the target location deviated from the cardinal axis by +/- 1, 2, or 3 pixels. In particular, we first obtained the posterior probabilities for scores estimated by fitting logistic regression models with the fitglm() Matlab function, and based on these scores, we calculated the AUC of the ROC curves using the perfcurve() function. The value of the AUC of the ROC curve could range between 0.5 and 1, with the former indicating a complete overlap between two distributions, and the latter indicating their complete separation.

Finally, we investigated whether the variability in the saccade initial directions was compensated online by the end of the saccades. To this end, we again replicated the analysis suggested by (Yoshida et al. 2008). To provide a sufficient amount of data points, we collapsed the trials across different pixel offsets within a saccade direction. In order to correct for the angular deviation introduced by the target pixel offsets in saccade trajectories, we rotated saccade trajectories about the point of origin by an angle by which the particular target offset deviated from the 0 pixel offset position of the corresponding direction and eccentricity. For example, if the saccade was 5 deg rightward and with +1 pixel offset, we rotated the saccade trajectory by an angle of 0.34 deg clockwise around the origin before pooling this condition with the 0 pixel offset condition; similarly, in the case of a 10 deg rightward saccades and −1 pixel offset, the rotation was by 0.17 deg anticlockwise.

Following (Yoshida et al. 2008), we then defined the initial direction as the vector angle from saccade start to the point on the saccade trajectory at which the amplitude reached 50% of the actual saccade amplitude. End direction was the same measure but based on the end point of the saccade. After that, we constructed histograms of the directional component (in 0.01 deg bins) of the end points and initial directions and convolved them with a Gaussian kernel with 1 deg s.d. We took the Full Width at Half Height (FWHH) of these smoothed distributions as a measure of their variability and computed the compensation index as 1 – (FWHH of the end points / FWHH of the initial directions). Thus, a compensation index larger than zero would indicate a compensation, and a compensation index equal to or less than zero would mean that the variability of initial directions was not reduced by the end of the saccades.

For the human data, we repeated the same essential analysis steps as for the monkeys. Thus, for each participant, we plotted the time courses of the average horizontal and vertical eye position traces for each orthogonal offset in the interval of −50 to 100 ms relative to the saccade onset, with the standard error of the mean as an estimate of the across-trial variability. Also, for each participant, we obtained two-dimensional visualizations of the time-courses of saccade trajectories. The quartile analysis was performed individually for each subject and each saccade direction but only 0 pixel offset conditions were taken into account in this case. Finally, when analyzing deviations introduced by the target orthogonal offsets in the course of the saccades, we also fitted GAMM’s but this time, we collapsed the data across participants and included a random smoother of time for a participant as an additional random effect in the model’s formula, in order to account for between-subject variability in the time courses of saccade trajectories. Thus, for each saccade direction, the fixed effects of the full model consisted of a categorical predictor of the target offset with seven levels of pixel offset (0, +/- 2, +/- 4, or +/- 8 pixels) and a factor-smooth interaction of the target offset and time, and the random effects included a by-session random smoother of time and a by-participant random smoother of time.

To relate the humans’ saccades to their perceptual reports, we also added a new eye movement analysis to the human data. In particular, the perceptual tasks had subjects classify the target location as being above or below the horizontal meridian (for horizontal saccades) or as being to the right or left of the vertical meridian (for vertical saccades). To compare whether perceptual biases in this target localization were related to oculomotor biases in eye movement landing positions, we calculated so-called oculometric curves (Kowler and McKee 1987; Stone and Krauzlis 2003). Specifically, for a given saccade (e.g. rightward), we measured the orthogonal component of the final landing position at saccade end (e.g. vertical eye position for a rightward saccade). We then classified the orthogonal landing position as indicating whether the eye classified the target as being on one side or the other of the visual meridian. For example, if the saccade was rightward and the vertical eye position at saccade end was >0, then we classified this trial as the saccade detecting the target as being “above the horizon”. On the other hand, if the vertical eye position at saccade end was <0, then this trial was one in which the saccade classified the target as being “below the horizon”. Across all trials and all pixel offsets, we could then take these classifications and plot them as psychometric curves, which we called the oculometric curves of the saccades (Kowler and McKee 1987; Stone and Krauzlis 2003). Based on these curves, we could calculate bias as the pixel offset shift that resulted in equal likelihood of saccadic classifications (e.g. 50% “above the horizon” saccadic responses). Because we found that cardinal saccades can have a small orthogonal offset in their landing positions after saccades (relative to orthogonal eye position at saccade onset), we performed the oculometric curve analysis using two reference frames. In one, we used the absolute 0 pixel offset position as the boundary for classifying saccades as being on one side or the other of 0. In the other, we used the median landing position in the 0 pixel condition as the reference 0 value based on which saccades were classifying a target as being above/below the horizon (for horizontal saccades) or to the right/left of vertical (for vertical saccades). In both cases, eye position at saccade onset for all saccades was re-aligned to 0 as in most of our other analyses.

### Behavioral report analyses in humans

First, for each participant and target direction, we calculated the proportions of ‘above’ and ‘right’ responses for horizontal and vertical targets, respectively, depending on the orthogonal target offset. We did this procedure both for the perceptual dual and perceptual control tasks. We then constructed psychometric curves and computed corresponding points of subjective equality (PSE’s) with the help of the Matlab toolbox for psychometric function estimation Psignifit 4, which uses Bayesian inference methods for constructing beta-binomial models of psychometric functions and has proven its robustness even for overdispersed data (Schutt et al. 2016). Finally, for each of the tasks, we obtained the means and their standard errors of PSE’s across participants. The oculometric curves were also fit with the help of the Psignifit 4 toolbox.

## Results

We tested the amplitude and directional accuracy of purely cardinal saccades of different sizes and directions. For the most precise characterization of eye movements, we used three highly trained rhesus macaque monkeys implanted with scleral search coils (Methods), which allow very high quality eye tracking. We first asked how often purely cardinal saccades can be observed after repetitions of tens or hundreds of saccades, and we then investigated the relationship between purely cardinal saccades and initial fixational eye position variability. To further demonstrate the remarkable directional accuracy of saccades, we also intentionally introduced minute deviations from pure cardinality in the target locations themselves (Methods), and we asked whether the saccades exhibited correspondingly small changes in trajectory to account for these deviations. Finally, we replicated all of these results in 14 naïve human subjects, albeit with a less reliable eye tracking technique (Methods). Our results demonstrate the impeccable quality with which saccades are generated, accounting for even tiny fixational eye position variability at their onset, and they also suggest that purely cardinal saccades may not be necessary in the first place, again by virtue of fixational eye position variability at saccade onset. This nulls a challenge of dealing with visual field discontinuities in the sensory and sensory-motor maps that are used to define saccade vector characteristics in retinocentric coordinates. These results have neurophysiological implications, in the SC and other structures, which will hopefully be researched in future investigations. In what follows, we first start with the rhesus macaque monkey observations, and we then describe our replication of these observations in human subjects.

### Purely cardinal saccades are extremely unlikely

To first illustrate our primary result, consider the example 10 deg rightward saccade shown in Fig. 1A from one monkey. The monkey was initially fixating a central fixation spot, which then jumped instantaneously by 10 deg rightward; there was no orthogonal component to the target jump (i.e. this was a 0 pixel offset condition from the task described in Methods). The visually-guided saccade that ensued was also 10 deg in size (see the horizontal eye position aligned on saccade onset in Fig. 1A). However, note that the vertical component of the saccade (light blue) still exhibited a clear intra-saccadic curvature of trajectory (magnified in the inset, and showing both an intra-saccadic oscillation as well as a final offset in eye position). Across all of our experiments and subjects, and with thousands of saccades, we found that such a small amount of curvature in purely cardinal saccades was obligatory, and practically never absent. Specifically, consider, once again, the example case of rightward saccades in Fig. 1A. We measured, for each saccade, the maximal eye position deviation in the vertical (orthogonal) direction during the saccade itself. We then measured the maximal eye position deviation in the same vertical (orthogonal) direction during a 100 ms pre-saccadic fixation interval for comparison. Across all saccades from this very same condition, we plotted cumulative distribution functions of both measures (Fig. 1B). The two distributions practically never overlapped: the maximal amount of deviation in vertical eye position during pre-saccadic fixation was small, and the 95^th^ percentile pre-saccadic deviation (0.072 deg) was still much smaller in size than at most the 1^st^ percentile deviation of intra-saccadic curvature in the second distribution (0.096 deg) (Fig. 1B). Therefore, purely “straight” rightward saccades are practically non-existent, at least in this monkey. This result is consistent with earlier observations of a so-called “cross-talk” between horizontal and vertical eye positions in cardinal saccades (Bahill and Stark 1977; King et al. 1986; Smit and Van Gisbergen 1990; Smit et al. 1990), and our goal here was to investigate whether such cross-talk was only due to noise, or whether it was functionally related to aspects like neural circuit discontinuities at the meridians and fixational eye position variability at saccade onset.

Even though we will return to analyzing results with human subjects again later on, and in more detail, Fig. 1C shows that the same phenomenon also holds true in human saccades. Specifically, the figure illustrates a case in which a human subject (tracked with a video-based eye tracker; Methods) made more than 1200 purely horizontal saccades of the same amplitude across many trials. Once again, practically all saccades were associated with a maximal vertical (i.e. orthogonal) component that was larger than any deviation in vertical eye position during pre-saccadic fixation. Therefore, both an example monkey (Fig. 1A, B) and an example human subject (Fig. 1C) demonstrate that purely cardinal saccades (in this case, horizontal) are essentially non-present (Bahill and Stark 1977).

It is important to note here that such a result is not trivially explained by a rotation of the reference frame for eye tracking relative to the subject’s head (whether with scleral search coils in monkeys or with a video-based eye tracker in humans). For example, if the magnetic fields of the scleral search coil system are not perfectly aligned with head orientation in the laboratory, then measured eye positions might exhibit cross-talk between horizontal and vertical channels, and a similar cross-talk can happen for humans because the video camera aimed at the human subjects’ eyes is below the level of eye sight (to avoid occluding the experimental display used to present stimuli). However, such a small amount of cross-talk is already corrected for with our calibration routines (Chen and Hafed 2013; Tian et al. 2016). Moreover, saccades in different cardinal directions exhibit idiosyncratic biases in their curvature (an example is shown later in Fig. 5C) that are not explained with a simple coordinate rotation. As we present in Discussion, other factors like translation of the entire eyeball in the orbit (Bahill and Stark 1977; Miller and Robins 1992; Oohira et al. 1983; Sylvestre and Cullen 1999) might still be at play, but our results in the remainder of this paper highlight an important additional contributor of fixational eye position variability at saccade onset that also needs to be considered.

**Figure 5.**
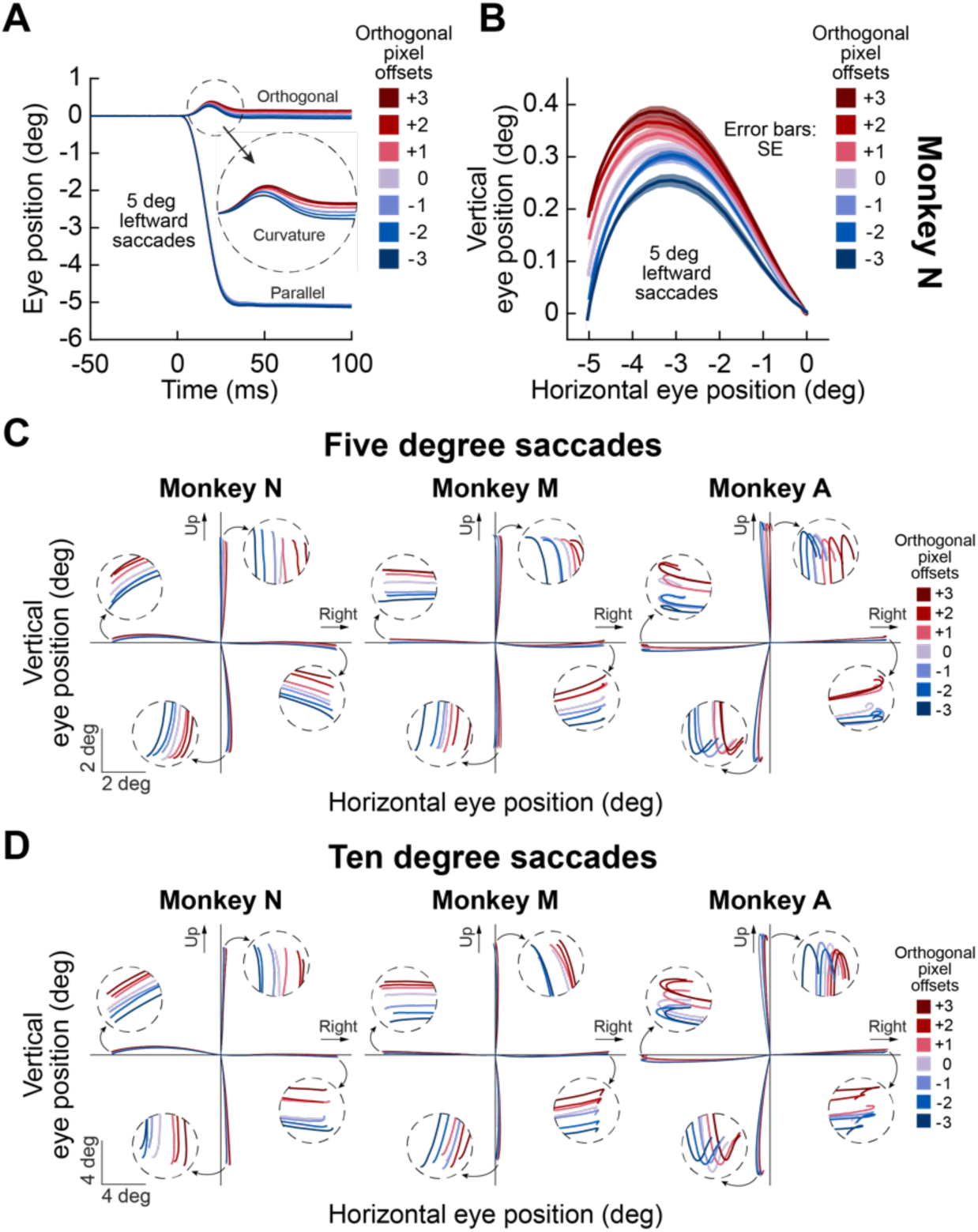
Cardinal saccade directions account for minute deviations of target positions from pure cardinality. **(A)** Average eye position traces from one monkey for an example saccade size and direction (5 deg leftward) when we offset the orthogonal component of target position jump from pure cardinality by a small amount (Methods). In the shown example, the target jumped by 5 deg to the left, and its vertical (orthogonal component) also jumped by 0 or +/- 1, +/- 2, or +/- 3 pixels (1 pixel is equivalent to 1.76 min arc). Despite the pixel jump being much smaller than the overall saccade size, there was a rank ordering of average saccade trajectories as a function of orthogonal pixel jump. We aligned all saccade starting points to zero in order to better visualize the influence of orthogonal pixel jumps. N = 87 to 101 saccades per pixel offset condition. **(B)** The same data as in **A**, but now showing the saccade trajectories in spatial coordinates. Upward pixel jumps (positive pixel values) deviated the saccades upward by a small amount relative to the 0 pixel condition, and downward pixel jumps deviated the saccades downward by a small amount. The range of average vertical landing positions across all pixel jumps was ∼12 min arc, which is remarkably close to the range of pixel jumps (7 pixels between −3 and +3; equivalent 12.32 min arc). Therefore, saccade directions accurately compensated their angular directions by as little as 0.34 deg from purely horizontal, despite a large angular radial amplitude of 5 deg. **(C)** Across all cardinal directions and monkeys, we aligned all saccade starting points to the origin (as in **A, B**), and we plotted the full trajectories of average eye position for each pixel offset. There was compensation for orthogonal pixel offsets in all cardinal directions; the insets magnify the final parts of eye trajectories, demonstrating a rank ordering of eye positions as a function of pixel offsets. N = 70 to 103 saccades per pixel offset and target direction in monkey N, 48 to 71 saccades in monkey M, and 20 to 49 saccades in monkey A. **(D)** Same as **C** but with 10 deg saccades. Similar conclusions could be made with respect to compensation for orthogonal pixel offsets. Despite the difference in amplitudes, all monkeys showed the same idiosyncratic curvature patterns as in **C**. N = 77 to 106 saccades per pixel offset and target direction in monkey N, 42 to 72 saccades in monkey M, and 31 to 49 saccades in monkey A.

**Figure 6.**
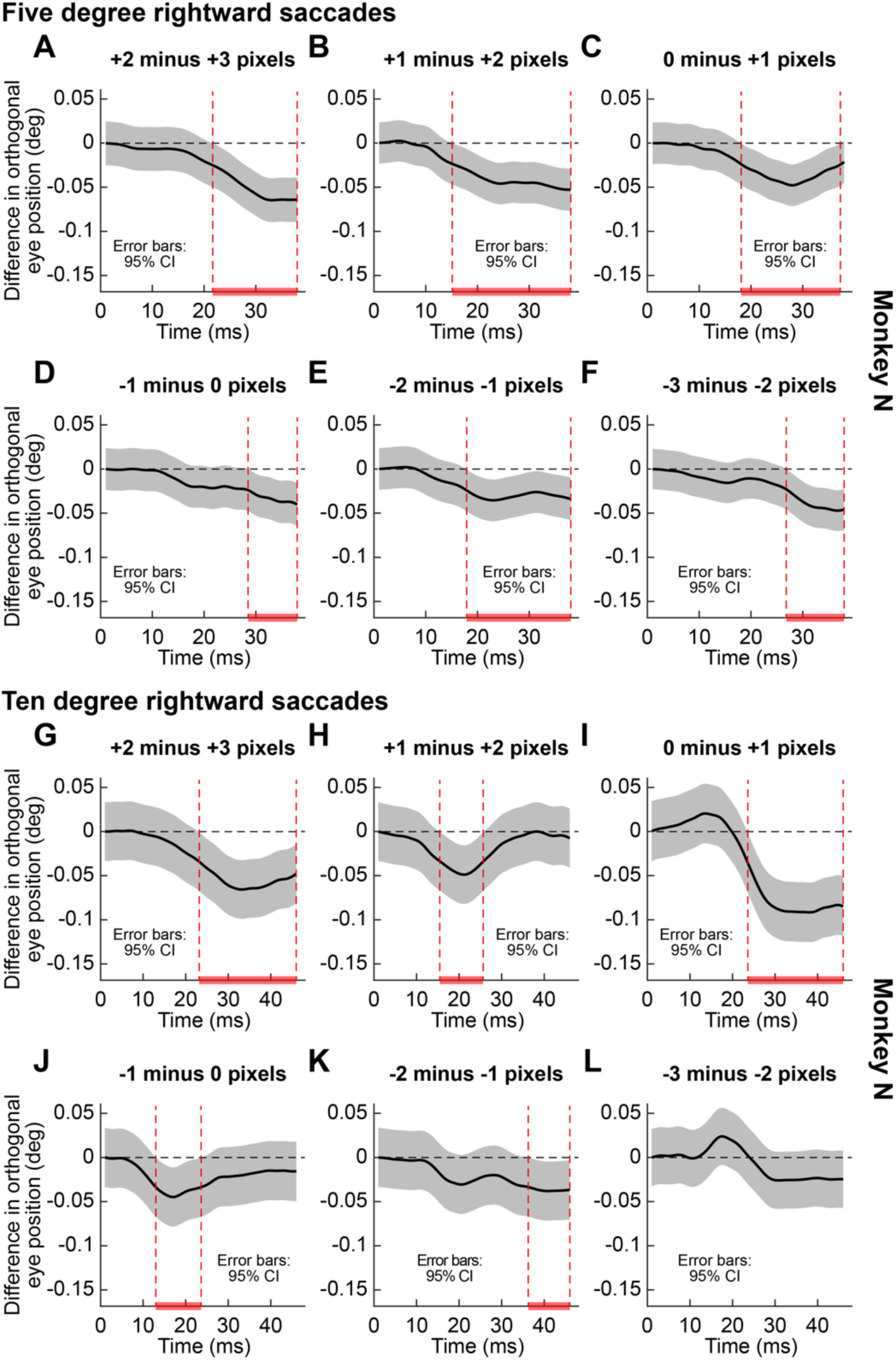
Saccades alter their trajectory, mid-flight, to curve towards the target when a small orthogonal component of target jump is introduced. **(A-F)** For an example monkey and saccade direction, we show the results of our generalized additive mixed model analyses (Methods) for statistically assessing the results of Fig. 5. For each 1-pixel difference between target position conditions, we evaluated whether saccade trajectories were different (i.e. whether saccade trajectories differentiated between minute changes in orthogonal component of target position jump). Each panel shows the modeled difference in orthogonal eye position between two conditions having a 1 pixel difference between target position jumps. The red bars on the x-axes indicate intervals of statistically significant differences in eye movement trajectories. In all cases, the eye movements deviated, intra-saccadically, to curve and more accurately foveate the slightly offset target. **(G-K)** Similar results for the larger 10 deg saccades. There was still intra-saccadic correction of saccade trajectories (only **L** shows a trend that did not reach significance at the 95% simultaneous confidence level). Monkey M showed very similar results (see text). For monkey A, the number of trials was too few to properly fit generalized additive mixed models (Methods).

**Figure 7.**
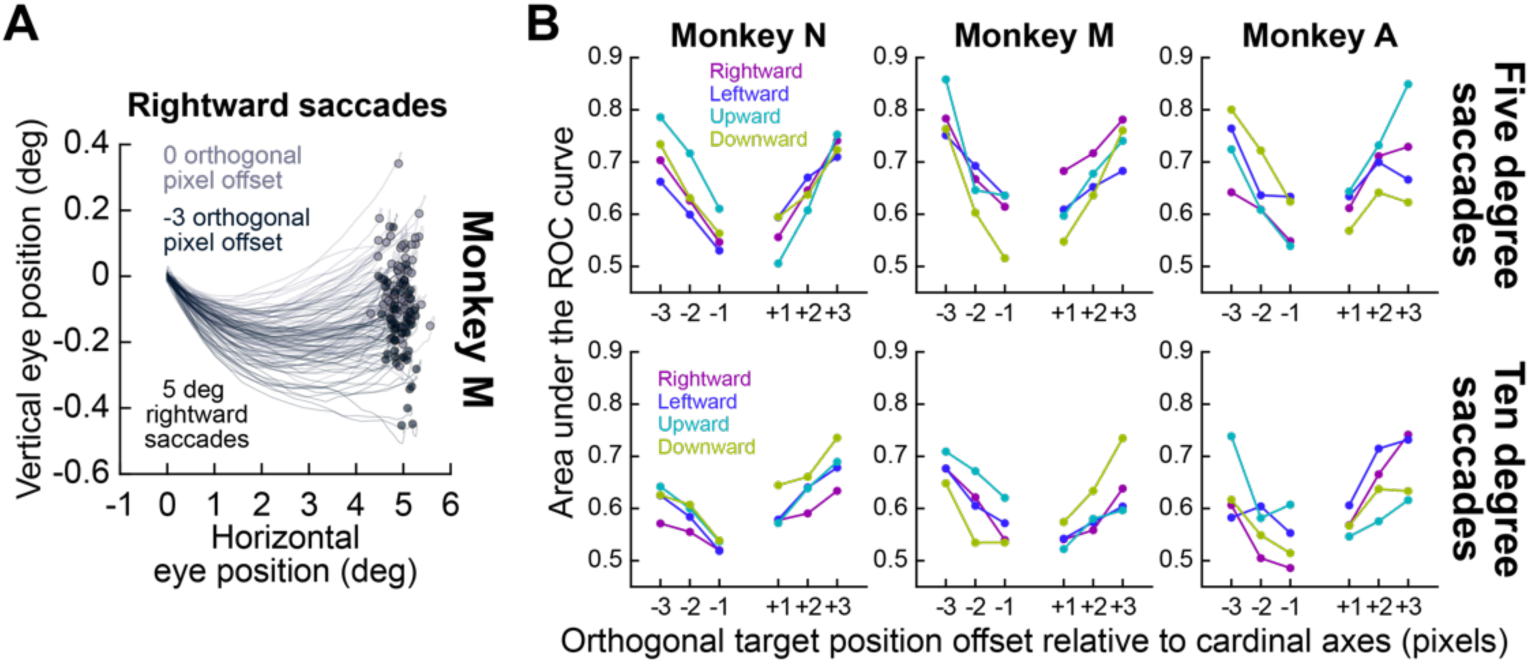
Discriminability of orthogonal pixel offset conditions across individual saccades. **(A)** Example eye trajectories (aligned to eye position at saccade onset) for two different pixel offset conditions. There was overlap in the saccade landing vertical (orthogonal) eye positions despite the corrections in average position seen in Figs. 5, 6. This prompted us to assess the discriminability of orthogonal landing eye position distributions in the different conditions. **(B)** For a given offset relative to the 0 pixel condition, we plotted distributions as in **A**, and we performed ROC analyses to assess how discriminable the orthogonal eye position distributions were. In all cases, even 1 pixel offsets had discriminability from 0 pixel offset that was above chance (>0.5), and discriminability expectedly increased with larger pixel offsets relative to 0 pixel offset. Discriminability was also greater than chance (>0.5) even for 10 deg saccades, for which saccade angular direction deviation needed to be as little as 0.17 deg (relative to a much larger radial angular amplitude of 10 deg). Note that for horizontal saccades, positive pixel offsets mean upward target position deviations from purely horizontal jumps, and for vertical saccades, positive pixel offsets mean rightward target position deviations from purely vertical jumps.

**Figure 8.**
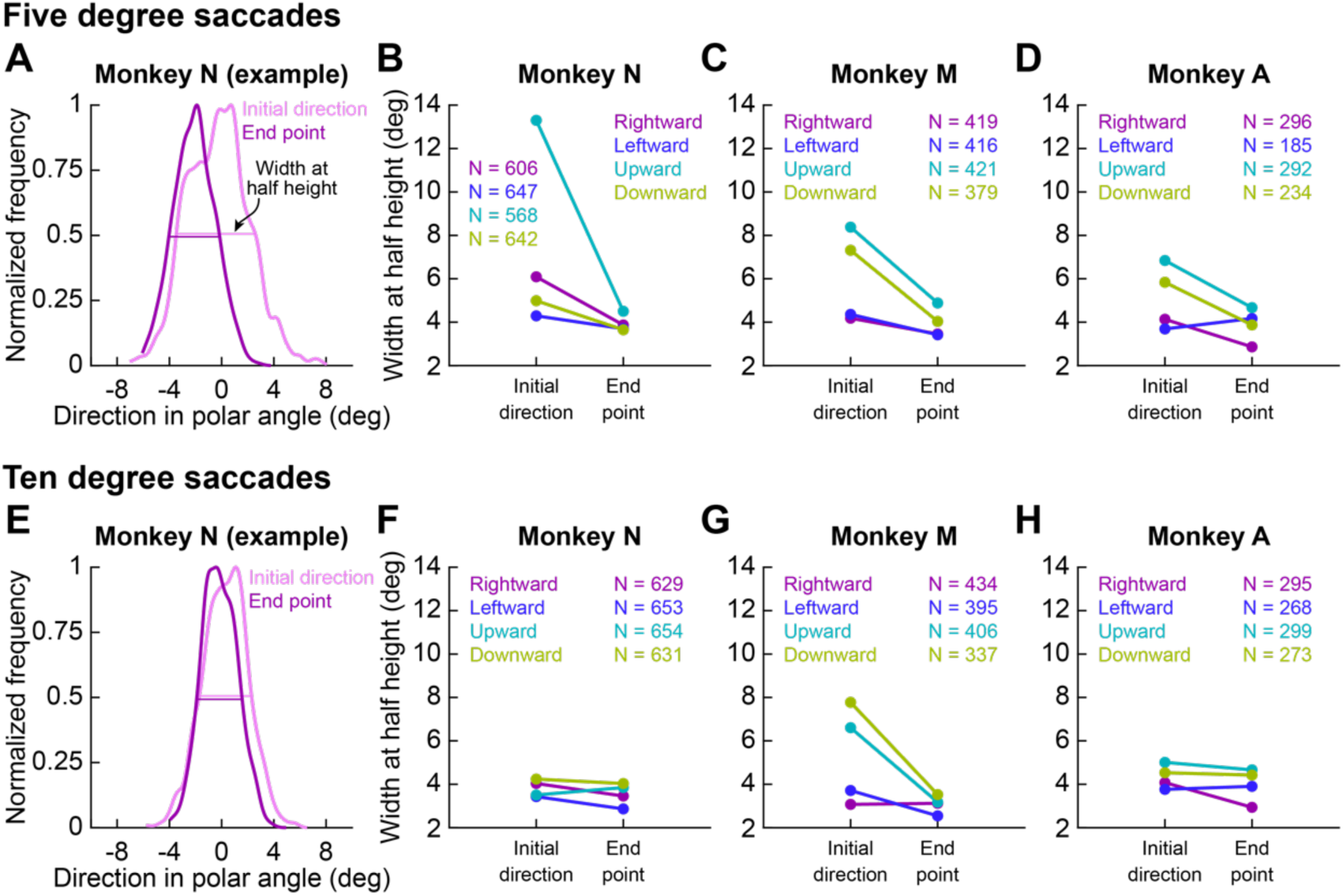
Intra-saccadic correction of saccadic curvature in purely cardinal saccades. **(A)** We adopted an analysis of saccade curvature similar to that in (Yoshida et al. 2008). For each saccade direction (in this example, 5 deg rightward in monkey N), we evaluated saccade curvature based on the first half of the trajectory (initial direction; light magenta) or based on the final end point (dark magenta). We obtained a distribution of angular directions of saccades in each case (Methods), and we assessed intra-saccadic correction by measuring the width of the distribution at half-height both mid-way through the saccade and by its completion (horizontal lines; the vertical level of the two lines was slightly jittered for the figure). In this example, there was a correction of saccade trajectory variability, such that the width of the distribution at half height was narrower by the end of the saccade than mid-way through it. **(B-D)** For each monkey and 5 deg saccades, there was a consistently lower half height width by saccade end than mid-way through the saccades (leftward saccades in monkey A were the exception). **(E-H)** Similar observations were made for 10 deg saccades (this time, the exceptions were upward saccades in monkey N, rightward saccades in monkey M, and leftward saccades in monkey A).

**Figure 9.**
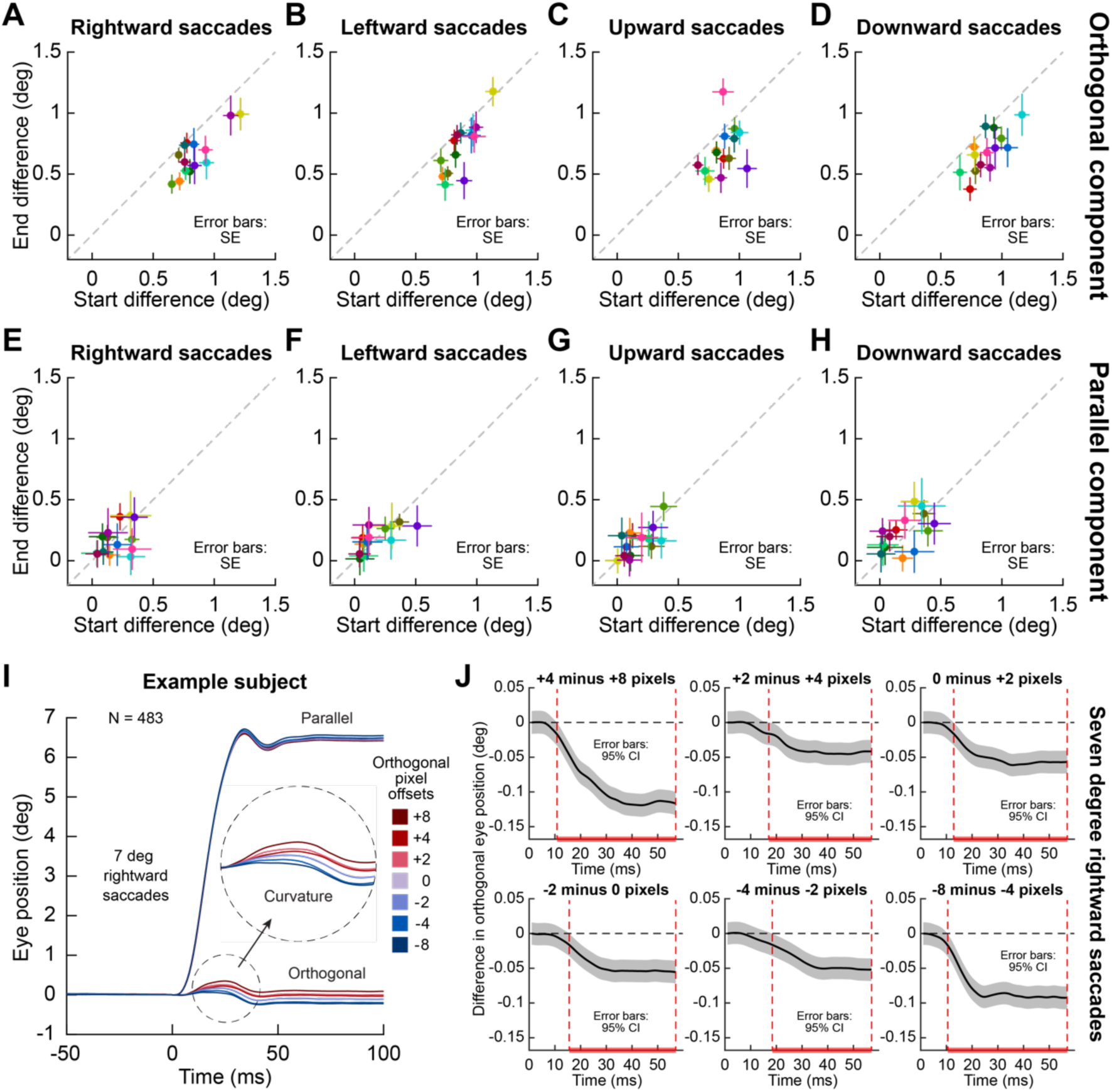
Similarity of effects in human saccades. **(A-D)** Human subjects performed a task with cardinal 7 deg saccades, and different orthogonal offsets like in the monkey experiments (Methods). We repeated the quartile analyses of Figs. 3, 4 for the 0 pixel condition (i.e. purely cardinal target jumps); each symbol shows results from one subject. In practically all subjects, saccade end point variability (along the orthogonal dimension) was smaller than initial end point variability. This suggests that human saccades also curve to correct for initial eye position at saccade onset due to fixational eye movements; this happens despite the use of video-based eye tracking, which is less ideal for properly measuring absolute fixational eye position. **(E-H)** Unlike in the monkeys (Figs. 3, 4), the parallel component of the saccades did not correct for the parallel component of initial fixational eye position. This could be due to the lower reliability of video-based eye tracking in accurately measuring fixational eye position, and it could also be due to the fact that video-based eye tracking measures the anterior segment of the eye, which oscillates significantly after saccade end (Nystrom et al. 2015; Nystrom et al. 2013). **(I)** With different orthogonal pixel offsets (1 pixel is 1.46 min arc), human saccades also curved accurately to account for the orthogonal target position deviation, just like in the monkeys (Figs. 5, 6). **(J)** Generalized additive mixed models of the human saccades, with the subject identity and session number as random effects in the models (Methods). Just like in the monkeys, human saccades corrected their trajectory, mid-flight, to more accurately foveate the targets in the different pixel conditions.

It is also important to note that there is, additionally, no biomechanical constraint of the eyeball that would make a result like in Fig. 1 trivially expected. For example, we electrically microstimulated the abducens nucleus (driving the lateral rectus eye muscle) in one monkey (in preliminary unpublished data), and we confirmed that the evoked lateral rectus muscle contractions were not associated with an intra-movement vertical curvature component to the measured eye position as we saw in Fig. 1A with real saccades; this seems to also be consistent with prior reports on abducens nucleus electrical microstimulation in the literature when the eye is in its primary position (Klier et al. 2006), although analyses of orthogonal eye position in abducens nucleus microstimulation studies is surprisingly rare (Anderson et al. 2009; Gandhi et al. 2008; Goldberg et al. 1998; Prsa et al. 2010).

Across all of our monkeys, we obtained consistent results for all cardinal saccade directions. Specifically, we repeated the analysis of Fig. 1 for either 5 deg or 10 deg saccades (all with the 0 pixel orthogonal offset in target position jump; Methods), and also for each of the four cardinal directions (right, left, up, and down). Monkey A had the least amount of trials collected, but even this monkey showed consistent results with the other two monkeys. This can be seen in Fig. 2, in which we plotted cumulative density functions separately for either horizontal or vertical saccades. As can be seen, the maximal deviation in orthogonal eye position component during a saccade was always consistently larger than the maximal orthogonal eye position deviation pre-saccadically. Also note that the distributions were more separated for the larger 10 deg saccades than for the smaller 5 deg saccades, likely due to the longer duration of the movements with 10 deg. Specifically, we performed one-sided two-sample K-S tests, and we confirmed that the orthogonal component of the 5 deg saccades was smaller than the orthogonal component of the 10 deg saccades for horizontal (monkey N: *D* = 0.492, p < 0.001; monkey M: *D* = 0.454, p < 0.001; monkey A: *D* = 0.5, p < 0.001) as well as vertical (monkey N: *D* = 0.382, p < 0.001; monkey M: *D* = 0.371, p < 0.001; monkey A: *D* = 0.53, p < 0.001) eye movements.

Therefore, the results of Figs. 1, 2 so far suggest that purely cardinal saccades are almost always associated with an orthogonal component to their trajectory (i.e. exhibiting a curvature and oscillation; Fig. 1A). This is consistent with early evidence from both humans and monkeys (Bahill and Stark 1977; Dodge 1917; Erkelens and Sloot 1995; King et al. 1986; Quaia et al. 2000; Smit and Van Gisbergen 1990; Viviani et al. 1977). Interestingly, from the perspective of the current study, such an orthogonal component suggests that it might not be necessary, in the first place, to ask whether purely cardinal saccades are possible at all, given visual field discontinuities at the meridians in retinocentric sensory and sensory-motor maps. In other words, purely cardinal saccades explicitly avoid pure cardinality by possessing a small amount of obligatory curvature in them. However, this does not, in any way, imply that saccades are simply sloppy eye movements in the vicinity of the visual field discontinuities. Rather, we think that the orthogonal component of saccades is there for a very good functional reason, as we describe next.

### Deviations from cardinality correct for initial fixational eye position variability

We next asked why purely cardinal saccades might still have an orthogonal component in their trajectory, which seems to be obligatory (Figs. 1, 2). Traditionally, saccade trajectory is analyzed primarily relative to eye position at the onset of the saccades. That is, all starting eye positions are re-referenced to be the origin of the analysis, and variability in landing eye positions is then characterized. This results in descriptions of landing error variability in a variety of contexts and under a variety of conditions, like with oblique versus cardinal saccades, and so on (Collewijn et al. 1988a; b; Greenwood et al. 2017; Quaia et al. 2000; van Beers 2007). However, in real life, the saccades start from an eye position that is inevitably variable across trial repetitions of the same saccade target jump, and this is due to incessant fixational eye position variability. Therefore, it may be the case that orthogonal components of saccade trajectory in purely cardinal saccades (e.g. Figs. 1, 2) are present in the saccades exactly to introduce a highly precise directional correction of saccade trajectory, which corrects for initial differences in the starting eye positions across trials.

Consider, for example, the sample situation shown in Fig. 3A, in which we analyzed all 5 deg rightward saccades from an example monkey (and from the 0 pixel orthogonal offset condition in target jump; Methods). Across trials, the initial starting eye position at saccade onset was expectedly variable due to fixational eye position variability. This is illustrated in the inset, in which we plotted the eye position at saccade onset across trials. We then divided the trials into four quartiles based on the orthogonal component of initial eye position relative to the intended horizontal saccade vector direction (i.e. since the saccade was horizontal, we divided initial eye position at saccade onset into four quartiles based on vertical eye position during fixation). When we plotted the saccade trajectories for the first and fourth quartiles in the main panel of Fig. 3A, we found that the saccades in the fourth quartile (starting from an upward fixational eye position relative to the first quartile) curved downward more than saccades in the first quartile, and this happened exactly to correct for the initial upward shift in fixational eye position at saccade onset. As a result, the vertical eye position after saccade end was mixed between the first and fourth quartile trials, when it was clearly segregated before saccade onset by default (Fig. 3A, main panel).

We then averaged the saccade trajectories of the four quartiles and plotted the saccade trajectories in Fig. 3B (left). The curves started with very different vertical components of fixational eye position (as per the design of the analysis), but they all converged together by saccade end. To obtain a quantitative measure of such convergence, we defined start and end differences in vertical (orthogonal) eye position between the first and fourth quartiles. As can be seen in Fig. 3B (left), the start difference in orthogonal eye position at the onset of the saccade (between the first and fourth quartiles) was relatively large (by definition of the quartile analysis). However, the end difference after the saccades was much smaller (Fig. 3B, left). Therefore, part of the significant curvature of purely cardinal saccades that we (and others) observed so robustly in Figs. 1, 2 is there in order to correct for initial deviations in starting eye positions at saccade onset. Note that such correction is remarkably precise. For example, for 5 deg saccades and a shift in orthogonal eye position between quartiles of approximately 0.1 deg due to fixational eye position variability (Fig. 3B, left), this means that 5 deg saccades can change their direction by as little as approximately 1.1 deg (in angular direction) in order to compensate for initial fixational eye position variability. For 10 deg saccades, this angular direction correction is even more precise (0.57 deg), but it was still present as we summarize later in Fig. 4.

Perhaps more remarkably, we also found corrections in saccade radial amplitudes as a function of initial fixational variability. Specifically, in Fig. 3B (right), we now split the trials into four quartiles again, but this time based on the parallel component of initial fixational eye position instead of the orthogonal component. The start difference in initial eye position at saccade onset between the first and fourth quartiles (Fig. 3B, right) was still larger than the end difference (Fig. 3B, right). Therefore, for an initial difference of approximately 0.1 deg between quartiles (due to fixational eye movements), saccades still corrected their amplitudes: for 5 deg saccades, this meant that a correction of amplitude by approximately only 2% of the total vector size was implemented (for 10 deg saccades, the correction was only 1%). Even though the directional correction (Fig. 3B, left) is more relevant for the visual field discontinuity question that is the primary topic of this study, we find it intriguing that saccade radial amplitudes can also be controlled so finely in order to correct for initial tiny fixational eye position variability (Fig. 3B, right). This suggests that saccades are not sloppy eye movements, but that they are under fine control even in the presence of fixational eye movements. Of course, there was still a small amount of global downward shift in the rightward saccades of Fig. 3, suggesting a small amount of bias in landing positions. We will assess such a bias later in Fig. 10 with respect to perceptual biases in target localization, but such a bias itself also suggests that purely cardinal saccades avoid neural circuit discontinuities in the representation of the visual meridians.

**Figure 10.**
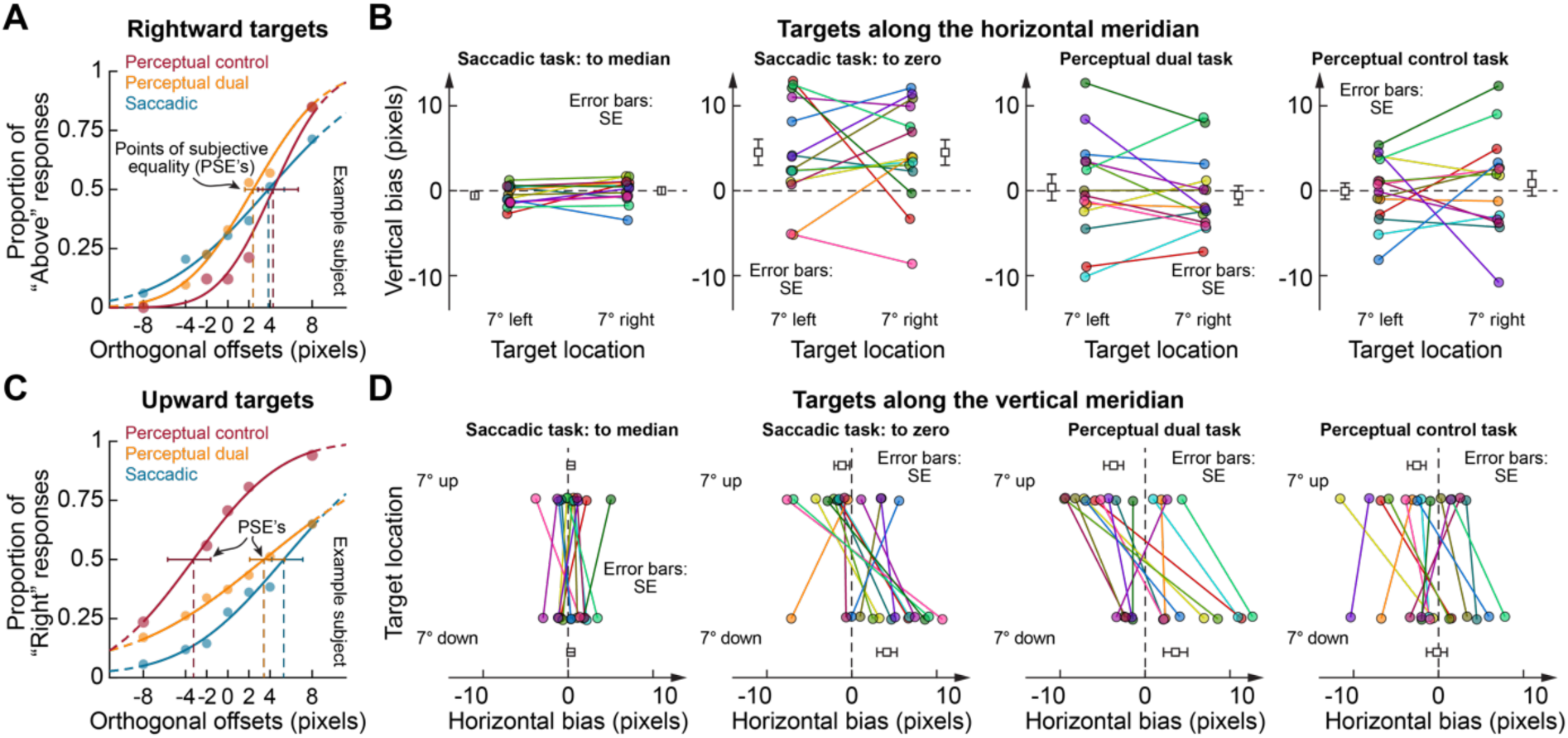
Different perceptual and oculomotor biases for small deviations of target positions from pure cardinality. **(A)** The main task of the human subjects was to generate a visually-guided saccade to a target (7 deg right, left, up, or down) that disappeared by saccade end (Methods). The orthogonal component of the target jump varied from pure cardinality by a small amount (1 pixel corresponds to 1.46 min arc). For horizontal saccades (rightward target shown as an example) the subjects judged whether the target was deviated from pure cardinality above or below fixation. The orange curve (“Perceptual dual”) shows an example subject’s psychometric curve. For the same subject, we judged the saccade end points (in the same task) and created an “oculometric curve” (labeled “Saccadic”) by finding whether the eye landed above or below the 0 pixel offset. In a control experiment (“Perceptual control”), the subjects maintained fixation, and the peripheral target was flashed for approximately 100 ms at the same locations as in the main task (Methods). From each curve, we estimated the point of subjective equality (PSE) as the orthogonal pixel offset resulting in equivocal perceptual reports or eye movement landing positions. **(B)** Leftmost panel: across subjects, we plotted PSE’s from oculometric curves like in **A** for horizontal saccades. In this case, we discriminated eye landing positions as being “above” or “below” based on the median landing position in the 0 pixel offset trials (landing positions above the median landing position in the 0 pixel condition were classified as “above” responses, and vice versa for landing positions below the median landing position in the 0 pixel condition). Each color is a subject; the black symbols show averages across the subjects. Across subjects, eye movements showed no bias relative to the median landing position of the control 0 pixel offset trials. Second to leftmost panel: same analyses but with biases in landing positions based on the absolute pure horizontal position of the 0 pixel offset condition (note that saccade starting points were all aligned in this analysis). Different subjects showed large biases. Rightmost two panels: psychometric curve biases from perceptual localization tasks. Perceptual biases were different from oculomotor biases. **(C)** Same as **A** but for upward saccades. **(D)** Same as **B** but for upward saccades. The directions of perceptual and oculometric biases in the dual task were the same.

Across all monkeys and all saccade directions and sizes, we also found consistent results with Fig. 3. In Fig. 4A-D, we plotted, in each panel, the end difference in orthogonal eye position between the first and fourth quartiles of trials (as in Fig. 3A) as a function of the start difference in orthogonal eye position, and we did this for 5 deg saccades. Figure 4A-D shows results similar to Fig. 3B (left). In each panel, we plotted each monkey’s data in one color, and for each monkey, the different symbols show analyses for all orthogonal pixel offset conditions in the behavioral task (Methods). That is, for each target position jump (whether it included pixel offsets in the orthogonal component or not), we checked whether end difference in orthogonal eye position compensated for start difference in orthogonal eye position. The large symbols show the average results of each monkey (from all orthogonal pixel offset conditions into one single grand average measure). As can be seen, for all monkeys and for all cardinal saccade directions, the end difference in orthogonal eye position after saccade end was consistently smaller than the start difference in orthogonal eye position before saccade onset (Fig. 4A-D). Only monkey A, with the least amount of trials (Methods), showed slightly larger variabilities in data tendencies. However, the overall results were all consistent. To statistically validate these observations, we computed IFCVI’s across the conditions (Methods). We found that the Monte Carlo probability of getting bootsrap IFVCI values larger than or equal to 1, which would indicate no compensation for initial eye position by the end of the saccade, was less than or equal to 0.014 in monkey N, less than or equal to 0.02 in monkey M, and less than or equal to 0.001 (except for leftward saccades; p = 0.172) in monkey A. Therefore, and as shown in Fig. 3B (left), a significant component of saccade curvature in purely cardinal saccades introduces directional trajectories in saccades, in order to correct for initial variability in (orthogonal) fixational eye position.

For the parallel component of initial fixational eye position variability (as shown in Fig. 3B, right), a similar conclusion was also reached (Fig. 4E-H). All three monkeys showed a correction in saccade radial amplitude to account for initial deviations in parallel eye position (either to be slightly closer to the peripheral saccade target along the parallel dimension to the saccade vector or slightly farther away from it). Statistically, the IFCVI’s revealed that the Monte Carlo probability of getting bootstrap IFVCI values larger than or equal to 1 (i.e. no compensation for initial eye position) was less than or equal to 0.007 except for leftward saccades (p = 0.224) in monkey N, less than or equal to 0.008 except for upward saccades (p = 0.135) in monkey M, and less than or equal to 0.018 in monkey A. This is remarkable, because it demonstrates that saccades are not sloppy in their radial amplitude. Rather, they can correct for radial amplitude changes that are as little as approximately 2% of the total desired saccade vector.

Perhaps more interestingly, all of these conclusions were still consistent for the larger 10 deg saccades (Fig. 4I-P). As can be seen, for practically all conditions and monkeys (Fig. 4I-L), the end difference in orthogonal eye position after 10 deg saccades was consistently smaller than the start difference. At the statistical level, the Monte Carlo probability of obtaining IFCVI’s of more than or equal to one (i.e. indicating no compensation) was less than or equal to 0.025 except for upward saccades (p = 0.379) in monkey N, less than or equal to 0.007 except for leftward (p = 0.65) and upward (p = 0.11) saccades in monkey M, and less than or equal to 0.025 in monkey A. This was also true for corrections of the parallel component of saccades; in this case, a 0.1 deg difference in initial fixational eye position at saccade onset (in the parallel direction; Fig. 3) is equivalent to a 1% correction in the amplitude of the 10 deg saccades. Yet, almost all conditions in Fig. 4M-P showed evidence for such small correction in saccade amplitude. The biggest exception was monkey M. At the statistical level, the probability of obtaining IFCVI’s more than or equal to one was less than or equal to 0.022 except for leftward saccades (p = 0.832) in monkey N and less than or equal to 0.003 except for rightward (p = 0.065) and downward (p = 0.928) saccades in monkey A; in monkey M, significant differences were revealed only for downward saccades (p = 0.001).

Therefore, slight changes in intra-saccadic trajectory, including curvature (Figs. 1, 2), are functionally relevant because they introduce compensatory components caused by variability in initial fixational eye positions at saccade onsets (Figs. 3, 4).

### Target-directed saccades are additionally sensitive to minute deviations of target positions from cardinality

If our interpretation from Figs. 3, 4 is correct, then we should also observe sensitivity in saccade trajectories if we experimentally introduce small deviations from pure cardinality in the saccade target position jumps. We thus analyzed trials from our conditions in which we explicitly introduced specific orthogonal pixel offsets in the target jumps (Methods). For example, in Fig. 5A, we had a monkey generate 5 deg leftward saccades. Across trials, when the target jumped by 5 deg to the left from central fixation, we additionally introduced a tiny orthogonal offset in the target position. The offset could consist of 0, +/-1, +/-2, or +/-3 pixels, with one pixel corresponding to 0.03 deg (1.76 min arc). That is, if the target had a 1 pixel offset in its jump, then the ideal saccade vector should have been 5 deg to the left horizontally and 1.76 min arc (0.03 deg) upward, and similarly for other pixel offset values. Remarkably, the saccades faithfully reflected these minute orthogonal target deviations from pure cardinality, consistent with our hypotheses and interpretations with respect to Figs. 3, 4. Specifically, in Fig. 5A, we plotted average horizontal and vertical eye position for each pixel offset condition. Note that in this figure, we re-referenced all saccades to start at 0 before averaging the eye positions. Given that fixation is accurate, on average, in monkeys (Tian et al. 2018; 2016), this is equivalent to taking the average fixational eye position (i.e. at the origin) and ignoring fixational eye position variability for the present analysis (such variability was already explicitly analyzed in Fig. 4). As can be seen, the vertical component of eye position during the horizontal saccades had a rank-ordering of eye positions that directly reflected the rank ordering of orthogonal pixel offsets (color codes in Fig. 5A). On the other hand, and as expected, the parallel component was much less scattered across the pixel offset conditions, since the parallel component of the target jump was always the same for all orthogonal pixel offset conditions. Therefore, not only do saccades correct for initial eye position deviations due to fixational eye position variability (Figs. 3, 4), they are also sensitive for targets with very small deviations from pure cardinality (of as little as 0.03 deg orthogonal component of the target jump).

The results of Fig. 5A can also be visualized in a different way, by plotting the two-dimensional trajectories of the saccades that were triggered. This is what we did in Fig. 5B. For all orthogonal pixel offset conditions, the horizontal component of the saccade was similar (5 deg leftward), as expected. However, the saccades deviated in their vertical dimension in a way that reflected, remarkably accurately, the orthogonal pixel offsets in the target position jumps. For example, there was a rank ordering of leftward saccade curves that was consistent with the rank ordering of the orthogonal target offsets (color codes in Fig. 5B). More interestingly, note how the final landing eye position across all pixel offsets had a total range of approximately 0.2 deg (compare the final vertical landing position for −3 pixel offset to the final vertical landing position for +3 pixel offset). This range was very close to the actual range of orthogonal target position deviations (a 7 pixel difference between −3 and +3 pixels corresponds to 0.2 deg in orthogonal target position deviation). Therefore, saccades faithfully deviate from pure cardinality when small deviations in target position from pure cardinality are introduced.

Across all monkeys and all cardinal directions, and also across our two saccade sizes (5 and 10 deg saccades), we made very similar observations. For example, in Fig. 5C, we plotted the two-dimensional average saccade trajectories for all pixel offsets in all three monkeys (for the 5 deg saccades). In all cases, the orthogonal component of the cardinal saccades was reflecting the orthogonal pixel offsets in target position jumps, as can be seen from the magnifications that we added to the figure (as insets) at each saccade end point. As can be seen, parallel landing positions for the different pixel offset conditions were similar (like in Fig. 5A, B), but orthogonal landing positions were different and reflected the orthogonal pixel offsets introduced by task design. Note also how in each monkey, upward and downward saccades both had a similar direction of small horizontal bias in their landing positions; as we stated earlier, this rules out a simple coordinate rotation as the explanations of Figs. 1, 2. Finally, Fig. 5D shows similar results with 10 deg saccades.

We next statistically confirmed that the saccades (whether they were 5 deg or 10 deg in size) did indeed correct their trajectories to foveate targets having small orthogonal position deviations from cardinality. To do this, we built generalized additive mixed models (GAMM’s) of the saccade trajectories (Methods). For each monkey, target eccentricity, and direction, the full models had lower AIC-values, and were hence a better fit, than the reduced models (Methods). Therefore, in each case, the best-fit models’ formula included a categorical predictor of the target pixel offset, a factor-smooth interaction of the target pixel offset and time, and a by-session random smoother of time; all estimated smooth functions showed non-linear trends, thus indicating the presence of curvature in saccade trajectories. The amount of variance explained by the models as measured by the adjusted coefficient of determination R^2^_(adj)_ was in the range of 0.312 to 0.778 in monkey N and in the range of 0.325 to 0.64 in monkey M. Using these models, we asked whether trajectories between pairs of neighboring pixel offsets deviated from each other in a statistically significant manner, suggesting sensitivity of the saccades to individual orthogonal pixel offsets. Almost all pairs of neighboring pixel offsets had statistically significantly different saccade trajectories. For example, Fig. 6 shows the results of GAMM analyses for an example monkey (N), for both 5 deg (Fig. 6A-F) and 10 deg (Fig. 6G-L) saccades. There was effectively always an intra-saccadic deviation of trajectories starting at approximately 15-27 ms after saccade onset. To be more specific, in monkey N, these differences were observed for 5 deg saccades, on average, starting from 18.753 ms +/- 5.712 ms s.d. at p < 0.05 for all adjacent pixel differences except for one out of 24 cases; for 10 deg saccades, these differences started at 19.165 ms +/- 6.755 ms s.d. on average and were observed at p < 0.05 for all adjacent pixel differences except for five cases (out of 24). For monkey M, the same GAMM analyses revealed that for 5 deg saccades, trajectories deviated between neighboring pixel pairs starting, on average, 15.346 ms +/- 4 ms s.d. after saccade onset at p < 0.05 (except for five out of 24 cases); for 10 deg saccades, these differences started, on average, at 19.04 ms +/- 7.099 ms s.d. and were observed at p < 0.05 for all neighboring pixel pairs (except for seven cases out of 24). Because monkey A had fewer trials, we could not run GAMM analyses on this monkey’s data (Methods), but we expect very similar results if more data were available to allow the models to converge (e.g. see Fig. 5C, D).

Therefore, the results of Figs. 1, 2 demonstrate that purely cardinal saccades have obligatory curvature in them (Bahill and Stark 1977), possibly to avoid visual field neural circuit discontinuities at the visual meridians. Moreover, the results of Figs. 3, 4 clarify that at least a component of cardinal saccade curvature is present to correct for variability in initial fixational eye position at saccade onset. And, finally, the analyses of Figs. 5, 6 allow us to conclude that saccades accurately foveate targets, even with tiny orthogonal offsets in their position, which require precise trajectory computations by saccades that are near pure cardinality.

We next asked whether landing variability, across trials, was also discriminable across pixel offsets. In other words, Figs. 5, 6 focused on average trajectories, so we next asked how discriminable the distributions of landing eye positions between pairs of pixel offsets were. For example, in Fig. 7A, it can be seen that the two shown orthogonal pixel offset conditions were associated with different average orthogonal landing positions (consistent with Fig. 5), but there was also overlap across repetitions. To evaluate this overlap, we performed ROC analyses on the orthogonal component of landing eye position for a given saccade direction (Methods). For example, for the example saccades in Fig. 7A, we evaluated the area under the ROC curve for discriminating the final vertical landing position of the saccades for each of the two pixel conditions. We found that even 1 pixel offsets (from 0) were associated with discriminable distributions of landing eye positions above chance (i.e. area under the ROC curve of >0.5). These results are illustrated in Fig. 7B for all monkeys. The x-axis indicates the pixel offset (relative to the 0 pixel condition), and the y-axis indicates the area under the ROC curve for discriminating a given pixel offset condition from 0 pixels. Different colors indicate different cardinal saccade directions. As can be seen, discriminability between distributions increased with increasing pixel offset sizes, as might be expected (and consistent with Fig. 5). Moreover, for 5 deg saccades, all areas under the ROC curve were >0.5 even for just 1 pixel offset. For 10 deg saccades, similar results were observed, but some conditions had weak discriminability for 1 pixel offsets, as might be expected.

Therefore, this is further evidence that saccades correct their trajectories to properly foveate peripheral targets. Naturally, this analysis re-referenced all saccades to start at 0, thus removing analysis of initial fixational eye position variability. This suggests that some of the remaining overlap between distributions in Fig. 7 could be explained by fixational eye position variability across trials (Fig. 4).

### Initial versus final saccade curvatures reveal mid-flight corrections to improve landing accuracy

Our analyses of Fig. 6 suggested that intra-saccadic trajectories may be modified, online, to improve saccadic landing accuracy. We next explicitly analyzed the variability in saccades early and late in the movements. We adopted an approach similar to that used in (Yoshida et al. 2008). Specifically, we measured saccade curvature halfway through the saccade as the angular direction of the vector connecting saccade onset to the orthogonal eye position deviation at the midpoint of the eye movement. This gave us a distribution of initial saccade directions (light magenta in Fig. 8A, E). We then computed the same direction measure at the end of the saccade (dark magenta in Fig. 8A, E). We found that saccades improved their precision by movement end, evidenced by a reduction in initial versus final curvature variability. This effect is illustrated in Fig. 8A for 5 deg saccades and in Fig. 8E for 10 deg saccades. The summaries of all monkeys and saccade direction are shown in the remaining panels of Fig. 8. Directional curvature distributions were almost always narrower by saccade end than mid-way through the saccades, likely helping to improve accurate landing on the targets (as seen in Fig. 5). To quantify these differences, we defined a compensation index as 1 – (FWHH of the end points / FWHH of the initial directions) so that a compensation index larger than zero would indicate a compensation of the initial variability by the end of the saccades (Methods; FWHH means Full Width at Half Height). We found that for 5 deg saccades, this index was in the range of 0.138-0.662 for all saccade directions in monkey N, 0.173-0.448 for all saccade directions in monkey M, and 0.308-0.337 for all saccade directions but leftward saccades (−0.131) in monkey A. For 10 deg saccades, the index was 0.048-0.165 for all saccade directions but upward saccades (−0.098) in monkey N, 0.315- 0.547 for all saccade directions but rightward saccades (−0.016) in monkey M, and 0.069-0.28 for all saccade directions but leftward saccades (−0.037) in monkey A. Overall, these results are consistent with previous reports that showed at least partial compensation of the initial direction errors by the end of the saccades (e.g. Aizawa and Wurtz 1998; Becker and Jurgens 1990; Erkelens and Sloot 1995; Erkelens and Vogels 1995; Quaia et al. 1999; Quaia et al. 2000; Yoshida et al. 2008).

Therefore, purely cardinal saccades exhibit online corrections and reductions in variability (Figs. 6, 8) to improve landing accuracy. All of this, in combination with Figs. 1-5, might suggest that an obligatory curvature in cardinal saccades might exist to avoid visual field neural circuit discontinuities, but that such curvature is functional and part of the control strategy for ensuring optimal eye movements.

### Humans show similar saccadic effects to monkeys, and their perceptual target localization is significantly different from their saccadic foveation

Finally, we were interested in confirming that human cardinal saccades show the same accuracy as monkey saccades, and we were also interested in relating saccadic localization of targets to perceptual reports of perceived target positions. We measured 7 deg saccades in 14 human subjects using a task similar to that used in the monkeys. Specifically, the target could jump by 7 deg in any one of the four cardinal directions. The orthogonal component of the target jump was either 0 pixels or it was +/-2, +/-4, or +/- 8 pixels (in this case, 1 pixel corresponded to 0.02 deg; Methods). The subjects performed a dual saccadic and perceptual task. Specifically, we detected saccade onset and removed the saccade target after saccade detection (target blanking occurred −14.48 ms +/- 9.59 ms s.d. relative to saccade end across all subjects and trials). After the saccades, the subjects had to report whether the target was above or below the horizontal meridian (for horizontal saccades) or whether it was to the right or left of the vertical meridian (for vertical saccades). Therefore, the saccadic landing positions provided us with an estimate of how saccades “localized” the peripheral targets, whereas the perceptual reports provided us with a measure of how the subjects perceived the targets relative to the horizontal and vertical meridians (Methods).

We first analyzed the saccade trajectories as we did with the monkeys. Even though the eye tracking technique was significantly worse than the scleral search coil method, and particularly for measuring initial fixational eye position variability, we still confirmed that human saccades showed the same properties as those that we reported above for the monkey saccades. For example, we already showed cumulative distribution analyses in Fig. 1C for an example subject with a large number of saccades, just to demonstrate that purely cardinal saccades (even with the 0 pixel offset condition) were extremely rare. The subjects explicitly collected in the present study allowed the same conclusions.

We then checked whether cardinal saccade curvature was related to the initial fixational eye position variability (as in Figs. 3, 4). To do this, we repeated the quartile analyses of Figs. 3, 4. We confirmed that saccade curvature compensated for variations in orthogonal eye position at saccade onset (Fig. 9A-D; each symbol represents an individual subject). The results were very similar to the monkey results. However, note that for corrections of the parallel components of saccades (Fig. 9E-H), the results were not as clear as in the monkeys. This likely reflects the effects of eye tracking technique. For example, video-based eye tracking is known to be affected by oscillations of the anterior segment of the eye by saccades (Nystrom et al. 2015; Nystrom et al. 2013), resulting in dynamic overshoots and oscillations (an example can be seen in Fig. 9I). As a result, it is harder to detect small changes in radial amplitude that are as little as 1% of the total saccade vector (7 deg), especially with the significantly worse performance of video-based eye trackers for classifying absolute fixational eye position variability when compared to scleral search coil techniques. In other words, the quartiles themselves were likely contaminated in the analyses with the video-based eye tracker. In any case, it is very likely that parallel component corrections should still exist in humans like in the monkeys as well, even when it is known that variability along the radial dimension of saccades is generally higher than along the orthogonal dimension (Greenwood et al. 2017; van Beers 2007; van Opstal and van Gisbergen 1989).

Finally, we checked for corrections in saccade trajectories with small orthogonal displacements of target position. This effect is shown clearly in Fig. 9I for a sample subject, replicating Fig. 5. As can be seen, the orthogonal component of eye position faithfully reflected the orthogonal component of target position jump, suggesting accurate saccade trajectory deviation. Indeed, GAMM analyses (with both by-subject and by-session random smoothers of time included as random variables) revealed corrections in saccade trajectory across all neighboring pairs of pixel offsets, like with the monkey eye movement analyses (Fig. 9J). In particular, these differences were observed, on average, starting from 13.59 ms +/- 2.53 ms s.d. at p < 0.05 for all adjacent pixel differences. Again, the models’ selection procedure (Methods) showed that the full models had better fits than the reduced models for each saccade direction; the amount of variance explained by the models as measured by R^2^_(adj)_ was in the range of 0.348 to 0.457. Therefore, we could conclude that human saccades show very similar effects to the monkey saccade effects that we presented in detail earlier.

Having established the human oculomotor effects, we were now interested in assessing perceptual localization. Specifically, and as stated above, the subjects (in addition to making a target-directed saccade) had to perceptually indicate whether they thought the peripheral target was deviating from pure cardinality (right or left of vertical for vertical saccades; or above or below the horizon for horizontal saccades). To avoid providing a foveal reference frame, we blanked the saccade target some time after saccade onset detection (Methods). We created psychometric curves of perceptual localization. For example, in Fig. 10A (“perceptual dual”), we plotted the fraction of trials in which the subject reported the peripheral horizontal target to be above the horizontal meridian. An expected psychometric curve was observed, which allowed us to obtain the perceptual bias (the pixel offset giving rise to 50% “above” reports). To check whether the dual nature of the task (saccade plus perceptual localization) might have caused the subjects to be perceptually biased, we also ran a control experiment in which the subjects maintained fixation, and the peripheral target was only flashed for 100 ms. The psychometric curve obtained was called “perceptual control” in Fig. 10A. Finally, we used eye movement end points to obtain an “oculometric curve” (Methods). If the vertical component of landing eye position was above zero, we designated the trial as revealing that the eye “signaled” an “above” response, and vice versa for landing eye position below zero. The resulting oculometric curve was labeled “saccadic” in Fig. 10A.

As can be seen, both perceptual and oculometric analyses revealed biases in peripheral target localization. However, there was no clear relation between perceptual and oculomotor effects. For example, for the same example subject in Fig. 10A, results with vertical saccades (Fig. 10C) showed very different biases. Across all target locations and subjects, we summarized the bias results in Fig. 10B (for horizontal targets) and in Fig. 10D (for vertical targets). For oculometric curves, we assessed bias (across subjects) in two ways (the leftmost two panels in Fig. 10B, D). In particular, because of slight biases in overall landing positions even with 0 pixel offset (e.g. Fig. 5C for monkeys), we first asked whether we could classify landing positions relative to the median landing position with the 0 pixel offset condition (leftmost panel in Fig. 10B, D). Expectedly from Figs. 3-6, oculometric curves with this approach showed very little eye movement classification bias across all subjects; in that regard, perceptual localization biases were much higher (rightmost two panels in Fig. 10B, D) than oculometric biases. The second oculometric analysis ignored the 0 reference of the eye movements with 0 pixels and just classified eye landing position as being either above or below pure 0 cardinality (in absolute display coordinates); now, oculometric curves based on the absolute 0 orthogonal position of the target (on the display) showed larger and more variable biases across subjects (second to leftmost panel in Fig. 10B, D). Interestingly, perceptual reports still did not match with oculometric reports in these analyses (rightmost two panels in Fig. 10B, D). The only exception might have been for the vertical targets, in which most subjects showed a stronger rightward bias in both perception (in the dual task) and saccades for downward targets when compared to upward targets.

Therefore, human cardinal saccades showed the same effects as the monkey cardinal saccades (Fig. 9). When we probed human perception in relation to the same saccades (Fig. 10), we found that subjective percepts of the vertical and horizontal meridians were generally not related to the eye movement performance. In fact, if the eye landing position with 0 pixel offsets is taken as the reference with which the oculomotor system defined the true meridian locations, then it can be concluded that eye movements were much less biased than perception in peripheral target locations (leftmost panels in Fig. 10B, D).

## Discussion

We studied the properties of purely cardinal saccades, motivated by the question of how neural circuit discontinuities at the visual meridians in sensory and sensory-motor maps may be handled. We found that purely cardinal saccades are practically non-existent, consistent with earlier classic observations (Bahill and Stark 1977; Dodge 1917; Erkelens and Sloot 1995; King et al. 1986; Quaia et al. 2000; Smit and Van Gisbergen 1990; Viviani et al. 1977). This is a way, in our view, to functionally avoid neural circuit discontinuities. However, such a feature of cardinal saccades does not mean that they are sloppy. Rather, their curvature compensates for initial fixational eye position variability at saccade onset, and it also allows redirecting of saccades to target positions that have very small orthogonal position deviations from pure cardinality. Finally, perceptually, it may be argued that subjective reports of the visual meridians are significantly worse than the reference frame of the oculomotor system for accurately directing saccades with sensitivity to individual pixel offsets in orthogonal target positions.

We find our results in relation to fixational eye position variability (e.g. Figs. 3, 4, 9) to be very interesting. There have been many intriguing hypotheses about the functional roles of fixational eye movements in the literature. For example, these movements were hypothesized to aid in avoiding perceptual fading due to stable images on the retina (Ditchburn et al. 1959; Ditchburn and Ginsborg 1952; Martinez-Conde et al. 2006; Nachmias 1961). Moreover, they are thought to influence cognitive processing in covert visual attention (Engbert and Kliegl 2003; Hafed 2013; Hafed et al. 2015; Hafed and Clark 2002; Laubrock et al. 2005; Tian et al. 2016). And, fixational eye movements can also optimize eye position (Ko et al. 2010; Poletti and Rucci 2013; Skinner et al. 2019; Tian et al. 2018), and they can alter the spatiotemporal profile of luminance information in images (Khademi et al. 2020; Kuang et al. 2012; Malevich et al. 2020; Rucci et al. 2007). Here, we identified another functional role for fixational eye movements: they avoid the need for purely cardinal saccades, which are computationally complicated by neural circuit discontinuities in the representation of the visual meridians.

Complementing the spatial uncertainty associated with fixational eye movements at saccade onset is an impeccable directional and amplitude ability of saccades. We found that saccades can compensate for orthogonal deviations from pure cardinality of as little as 0.03 deg or 1.8 min arc. This amount is almost as small as the diameter of an individual foveal cone photoreceptor (Borwein et al. 1980), and, yet, the eye movement system is sensitive enough to directionally deviate an eye movement that is at least two orders of magnitude larger in size in order to ensure proper foveation. This represents a remarkable control property of the oculomotor system for saccades, and it is not in line with some ideas that saccades may be sloppy in their landing scatter in general. Rather, we found that saccades started improving their trajectory control as early as approximately 15 ms after saccade onset (Figs. 6, 9J). Moreover, our ROC analyses revealed discriminability >0.5 even for 1 pixel offset conditions, and at least in one analysis in Fig. 10, oculometric discrimination of target positions was much less biased than subjective perceptual localization (Fig. 10B, D; leftmost panels). This is also consistent with existing literature showing that saccades can exhibit better performance than different measures of perceptual localization (Greenwood et al. 2017; Vishwanath and Kowler 2003). And, on top of that, our results of high sensitivity of saccades to tiny orthogonal target position deviations were not in any way forced by the task demands themselves. For example, even though our monkeys were well trained, the pixel offsets that we used were negligible in size relative to the virtual computer windows implemented for rewarding the animals at trial ends.

We are also intrigued by the idea that saccadic curvature with intra-saccadic oscillations (Aizawa and Wurtz 1998; Bahill and Stark 1977; Dodge 1917; Erkelens and Sloot 1995; Erkelens and Vogels 1995; King et al. 1986; Quaia et al. 1999; Quaia et al. 2000; Smit and Van Gisbergen 1990; Viviani et al. 1977; Yoshida et al. 2008) seems to be intrinsic to all saccades, even for so-called “straight” purely cardinal movements. As we stated earlier, we do not think that this observation is explained by just a rotation of reference frames (relative to the magnetic fields in the scleral search coil technique or relative to the camera orientation in video-based eye tracking). Also, such curvature is not likely to be fully explained by a biomechanical limitation of the eyeball. In fact, the medial rectus and lateral rectus extraocular muscles should not cause an orthogonal oscillation of the eyeball as they pull the eye horizontally from its primary position, and they should also not cause a cross-talk for other eyeball movement axes. Therefore, at least for horizontal eyeball rotations from the primary position, cross-talk is not necessarily to be expected from a biomechanical perspective. Having said that, large deviations in vertical eye position from the primary position might be associated with torsional components related to Listing’s law (Klier et al. 2006). It remains to be seen whether large vertical saccades (e.g. 10 deg) would be associated with a torsional component that can be registered during eye tracking as an orthogonal eye movement component. At the very least, with our video-based eye tracker, this is not likely to be the case, because our eye tracker is not sensitive to torsion. Other factors that can cause cross-talk between horizontal and vertical eye position measurements would include retraction of the eyeball in the orbit during saccades (Bahill and Stark 1977; Miller and Robins 1992; Oohira et al. 1983; Sylvestre and Cullen 1999) or vergence eye movements in association with vertical saccades (Enright 1989). It remains to be seen whether such additional factors are also sensitive to the small pixel offsets that we used in our studies or not.

In any case, all of these questions should be tested in much more detail in the future. For example, electrical microstimulation of the abducens and oculomotor nuclei, which innervate the lateral and medial rectus muscles, respectively, could be used to cause contractions of individual eye muscles while carefully measuring all components of eyeball rotations. As stated above, we have ongoing unpublished preliminary experiments in this direction, in which we activate the lateral rectus muscle but without observing a cross-talk in vertical eye position like the one that we have seen with real horizontal saccades. Therefore, an important future neurophysiological direction should be to follow up on these observations, and also to investigate the issue of intra-saccadic trajectory oscillations (Ghasia and Shaikh 2014; Ramat et al. 2005; Ramat et al. 2008; Yee et al. 1994) in much more detail. Indeed, it is surprisingly hard to find literature on abducens or oculomotor nucleus microstimulation with reports of vertical eye position measurements, and this is an important area of future research on the saccadic system. The net result could be that descriptions of saccadic trajectory variations as reflecting only noise (at least in the initial half of saccades) (Abrams et al. 1989; Harris and Wolpert 2006; 1998; Quaia et al. 2000) might be revised to reveal much more precise oculomotor control. Based on previous work, one might even anticipate that every spike in the efferent pathway from SC to eye muscles can have a deterministic impact on trajectory. For example, there is new evidence that even single spikes in baseline ongoing SC activity in individual neurons are sufficient to modulate the kinematics of individual saccades if the spikes occur intra-saccadically (Buonocore et al. 2020).

Finally, also at the neurophysiological level, it is important to consider the fine structure of RF’s near visual field discontinuities in retinocentric maps. For example, as stated in the Introduction, in the SC, it is suggested that the vertical meridian is represented by both SC’s. However, again here, it is surprisingly hard to find literature on detailed individual RF mappings near the vertical and horizontal meridians. Based on our own experience, we find that RF’s near the vertical meridian seem to still have preference for the contralateral side even when their RF’s extend spatially in the ipsilateral hemifield (Chen et al. 2019; Hafed and Chen 2016). That is, the RF hotspot location is still lateralized even when the RF extent does cross the vertical meridian discontinuity. A similar problem might occur for the horizontal meridian, since the SC also has a functional discontinuity at this meridian (Hafed and Chen 2016). An intriguing possibility is that the fine scale structure of SC RF’s near the visual meridians might explicitly avoid representation of pure cardinal directions in individual neurons, such that purely cardinal saccades are not possible. An implication of this is that RF’s near the visual meridians (both horizontal and vertical) might be patchy in their fine scale structure, and another implication of this is that purely cardinal saccades might exhibit intra-saccadic oscillations (e.g. Fig. 1A): the neural activity controlling saccade trajectories could keep oscillating between the two sides of the unrepresented discontinuity. Moreover, a small bias in landing position (e.g. Figs. 5, 10) might additionally be expected. This and related questions warrant much deeper investigation in the future, and they would add to extensive evidence that the landscape of SC activity plays an important role in saccadic curvature, whether this curvature occurs naturally (Port and Wurtz 2003), as a result of SC microstimulation (Noto and Gnadt 2009), or in tasks involving distractors or attention away from the saccade target (Laidlaw and Kingstone 2010; McPeek et al. 2003; McPeek and Keller 2001; McPeek et al. 2000; McSorley et al. 2006; van Leeuwen and Belopolsky 2018; Walker et al. 2006; White et al. 2012).

## Acknowledgements

We were funded by the Werner Reichardt Centre for Integrative Neuroscience (CIN), which was originally created through excellence cluster funds (EXC307) from the Deutsche Forschungsgemeinschaft (DFG). The project was supported by CIN intramural grant number Mini-GK 2017-04.

